# Behavior-dependent spatial maps enable efficient theta phase coding

**DOI:** 10.1101/2021.04.14.439787

**Authors:** Eloy Parra-Barrero, Kamran Diba, Sen Cheng

## Abstract

Navigation through space involves learning and representing relationships between past, current and future locations. In mammals, this might rely on the hippocampal theta phase code, where in each cycle of the theta oscillation, spatial representations start behind the animal’s location and then sweep forward. However, the exact relationship between phase and represented and true positions remains unclear. Developing a quantitative framework for the theta phase code, we formalize two previous notions: in spatial sweeps, different phases of theta encode positions at fixed distances behind or ahead of the animal, whereas in temporal sweeps, they encode positions reached at fixed time intervals into the past or future. These two schemes predict very different position representations during theta depending on the animal’s running speed. Paradoxically, in some studies sweep length has been shown to increase proportionally with running speed, consistent with temporal sweeps, whereas in other studies individual place field parameters such as field size and phase precession slope were shown to remain constant with speed, consistent with spatial sweeps. Here, we introduce a third option: behavior-dependent sweeps, according to which sweep length and place field properties vary across the environment depending on the running speed characteristic of each location. Analyzing single-cell and population variables in parallel in recordings from rat CA1 place cells and comparing them to model simulations, we show that behavior-dependent sweeps uniquely account for all relevant variables. This coding scheme combines features and advantages of both spatial and temporal sweeps, revealing an efficient hippocampal code.

**Significance:** To learn the structure of the world and the consequences of our actions, information about the past must be carried through to the present and linked to what is currently happening. To plan, desired future states and the predicted outcomes of actions must be represented. In mammals, including humans, hippocampal neurons are thought to encode such representations of past, present and future states at different phases of the theta oscillation. However, the precise hippocampal phase code remains unknown. We show that two previous ideas are incompatible with each other and with rat experimental data. So, we propose a new coding scheme that synthesizes features from both ideas and accounts for all relevant observations.

## Introduction

Hippocampal place cells elevate their firing rates within circumscribed regions in the environment (“place fields”) (O’Keefe and Dostrovsky, 1971). In addition to this rate code, place cells display phase coding with respect to ~8 Hz theta oscillations of the hippocampal local field potential (LFP) (Vanderwolf, 1969). As animals cross a cell’s place field, the cell fires at progressively earlier phases of the oscillation, a phenomenon known as theta phase precession (O’Keefe and Recce, 1993). The importance of phase precession becomes clearer at the population level. Within each cycle of the theta oscillation, the first cells to fire typically have place fields centered behind the current position of the animal, followed by cells with place fields centered progressively ahead, generating so-called theta sequences (*SI Appendix*, Fig. S1) (Skaggs et al., 1996; Dragoi and Buzsáki, 2006; Foster and Wilson, 2007; Maurer et al., 2012; Gupta et al., 2012; Muessig et al., 2019). Thus, the position *r*(*t*) represented by the hippocampal population at time *t* deviates from the physical location *x*(*t*) of the animal, sweeping from past to future positions during each theta cycle. This phenomenon appears to extend beyond just the representation of spatial positions in the hippocampus to encompass the representation of past, present and future events more generally (Dragoi and Buzsáki, 2006; Cei et al., 2014; Terada et al., 2017). In support of this, theta phase precession or theta sequences have been reported for cells responding to elapsed time (Pastalkova et al., 2008; Shimbo et al., 2021), or particular events or behaviors such as jumping (Lenck-Santini et al., 2008), pulling a lever, or sampling an object (Terada et al., 2017; Aronov et al., 2017; Robinson et al., 2017), and has been observed in structures beyond the hippocampus (Kim et al., 2012; Hafting et al., 2008; Jones and Wilson, 2005; van der Meer and Redish, 2011; Tingley et al., 2018; Tang et al., 2021). These findings suggest that the theta phase code plays a role in supporting a wide array of cognitive functions, such as sequential learning (Lisman and Idiart, 1995; Skaggs et al., 1996; Reifenstein et al., 2021), prediction (Lisman and Redish, 2009; Kay et al., 2020) and planning (Johnson and Redish, 2007; Erdem and Hasselmo, 2012; Bolding et al., 2020; Bush et al., 2015).

However, for the hippocampal theta code to be useful, there must be a consistent mathematical relationship between theta phase, represented position *r*(*t*), and the animal’s true location *x*(*t*). Surprisingly, this relationship has not previously been made explicit. It is often stated interchangeably that activity at certain phases of theta reflect positions ‘behind’ or ‘ahead’ of the animal, or, alternatively, in its ‘past’ or ‘future’, but these statements hint at two fundamentally different coding schemes.

The first possibility is that different theta phases encode positions at certain fixed distances behind or ahead of the animal’s current location. For example, within each theta cycle, the represented position *r*(*t*) could sweep from the position 2 meters behind of the animal to the position 2 meters ahead. The hippocampus would thus represent positions shifted *in space*, which is why we refer to this coding scheme as the ‘spatial sweep’. Another possibility is that the hippocampus represents positions that were or will be reached at some *time* intervals into the past or future, respectively. We call this scheme the ‘temporal sweep’. For example, *r*(*t*) might start in the position that was reached 5 seconds ago and extend to the position projected to be reached in 5 seconds into the future. Walking at home, the position reached in 5 seconds might be a couple of meters ahead, whereas while driving in the highway, that position could be more than a hundred meters away. Thus, in temporal sweeps, the look-behind and -ahead distances adapt to the animal’s speed of travel, which could make for a more efficient code.

In the following, we set out to develop a quantitative description of the hippocampal theta phase code. First, we formalize the spatial and temporal sweep models. Intriguingly, as we discuss in detail below, different experimental results appear to support different models. The length of theta sweeps has been shown to increase proportionally with running speed, consistent with temporal sweeps (Maurer et al., 2012; Gupta et al., 2012). On the other hand, place field parameters, such as place field size or phase precession slope, do not change with speed, consistent with spatial sweeps (Huxter et al., 2003; Maurer et al., 2012; Schmidt et al., 2009; Geisler et al., 2007). Hence, we propose an additional coding scheme, the ‘behavior-dependent sweep’ that could potentially reconcile these apparently contradictory findings. According to this model, the extent of the theta sweep varies across the environment in proportion to the animal’s characteristic running speed at each location. Next, we analyze recordings from rat CA1 place cells and reproduce, in the same dataset, the paradoxical results just mentioned. Crucially, however, we find that place fields are larger and have shallower phase precession slopes at locations where animals typically run faster, consistent with the behavior-dependent sweep model. Finally, we compare simulated data generated from the three models based on experimentally measured trajectories and theta oscillations, and confirm that the behavior-dependent sweep uniquely accounts for the combination of experimental findings at the population and single-cell levels.

## Results

### Formalizing the theta phase code

It is generally held that the theta phase of a place cell’s spike reflects the distance traveled through the cell’s place field (Huxter et al., 2003; Geisler et al., 2007; Cei et al., 2014) or some measure of the distance to the field’s preferred location (Jeewajee et al., 2014; Huxter et al., 2008; Drieu and Zugaro, 2019). In a uniform population of such phase coding place cells, the represented position, *r*(*t*), sweeps forward within each theta cycle, starting at some distance behind the current position of the animal and ending at some distance ahead. We formalize such spatial sweeps in this equation:

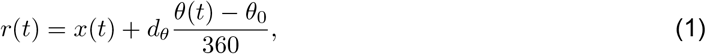

where *d_θ_* stands for the extent of the spatial sweep, which we call the theta distance, *θ*(*t*) is the current theta phase in degrees, and *θ*_0_ is the phase at which the population represents the actual position of the animal. *θ*_0_ controls the proportion of the sweep that lies behind vs ahead, although none of our results or analyses depends on the particular value of this variable.

At the single-cell level, the spatial sweep model predicts that place field size and the slope of the spikes’ phase vs. position relationship (phase precession slope) remain unaffected by running speed (Fig. 1B). This has been supported by several experimental observations (Huxter et al., 2003; Maurer et al., 2012; Schmidt et al., 2009; Geisler et al., 2007). However, the model does not account for results at the population level regarding theta trajectory length, defined as the difference between the maximum and minimum positions that *r*(*t*) represents within a theta cycle. Using a first order approximation, i.e., assuming that effects from acceleration and higher order derivatives are negligible, the theta trajectory length is *l* = *vT* + *d_θ_*, where *v* is the constant running speed, and *T* is the duration of a theta cycle (see Methods). Therefore, the model predicts that theta trajectory length increases only slightly with running speed due to the term *vT*, which represents the small change in *x*(*t*) due to the animal’s motion during the theta cycle (Fig. 1A, see also Fig. 1 in Maurer et al. (2012)). Contrary to this prediction, experimental results show that the theta trajectory length is roughly proportional to running speed (Maurer et al., 2012; Gupta et al., 2012).

**Figure 1:**
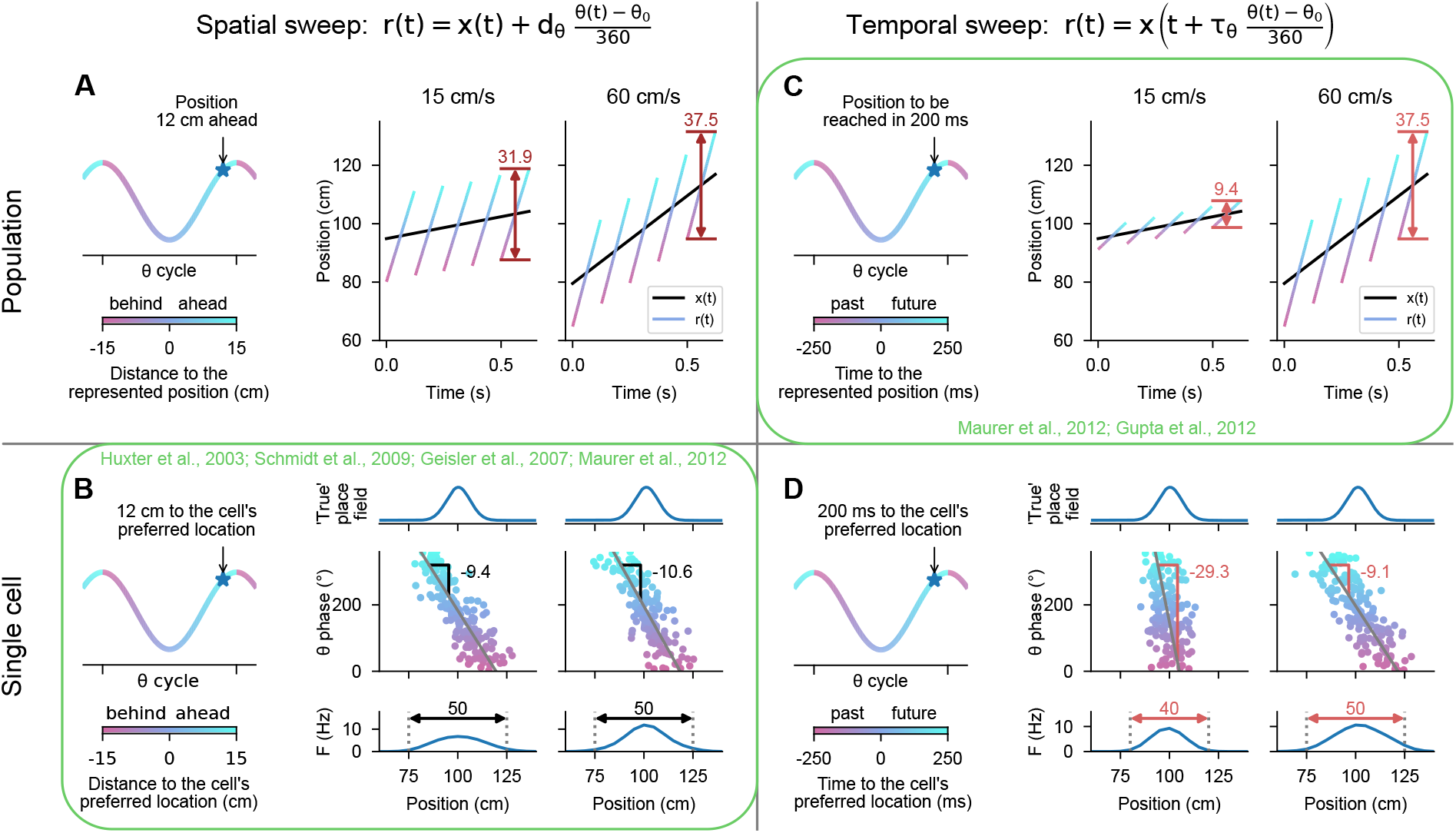
Effect of running speed on population and single-cell properties in the spatial and temporal sweep models. **A**: Left. At different phases of theta, the population represents positions shifted behind or ahead in space by *fixed distances*. Right. The black lines represents the rat’s actual position *x*(*t*) as it runs through a linear track; the color-coded lines indicate theta trajectories represented by the place cell population *r*(*t*). Since each theta trajectory finishes a fixed distance ahead of the current position, the length of the theta trajectories increases slightly with running speed in order to compensate for the change in the actual position of the animal during the theta cycle (37.5 vs. 31.9 cm). **B**: Left. At the single single cell level, the phases at which a cell spikes reflect the *distances* to the cell’s preferred location. Right. The cell’s preferred location is defined by its underlying ‘true’ place field (top). The cell fires proportionally to the activation of its true place field at *r*(*t*), generating a phase precession cloud (middle) and measured place field (bottom). Phase precession slopes and place field sizes remain constant with running speed. **C**: Left. At different phases of theta, the population represents the positions that were or will be reached at *fixed time intervals* in the past or future, respectively. Right. A higher running speed leads to a proportionally increased theta trajectory length since, e.g., the position that will be reached in 200 ms is further ahead in space at a higher than at a lower speed. **D**: Left. At the single-cell level, the phase of theta reflects the *time* to reach the cell’s preferred location. Right. At higher speed, the phase precession slope becomes shallower (−9.1 vs. −29.3 °/cm) and the size of the measured place field increases (50 vs. 40 cm) since, e.g., the cell will start signaling the arrival at the cell’s preferred location in 200 ms from an earlier position in space. Previous studies paradoxically support the spatial sweep at the single-cell level **B**, and the temporal sweep at the population level **C**.

An alternative sketched above is the temporal sweep model, where different phases of theta represent the positions which the animals had reached or will reach at some time intervals in the past and future, respectively (see also Itskov et al. (2008)). More formally,

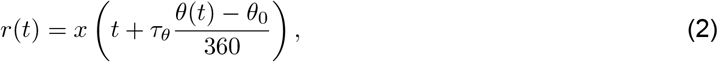

where *τ_θ_* stands for the extent of the temporal sweep. Again using a first order approximation, we derived that the theta trajectory length is proportional to running speed: *l* = *v*(*τ_θ_* + *T*) (Methods) (Fig. 1C). This is almost exactly the result reported by Maurer et al. (2012); Gupta et al. (2012). However, at the single-cell level, the temporal sweep model predicts that place field sizes and phase precession slopes increase with running speed (Fig. 1D) – in clear conflict with the experimental results mentioned above.

These conflicting results originate almost exclusively from different studies, raising the possibility that differences in experimental procedures or analytical methods caused the apparent contradiction. If so, either spatial or temporal sweeps, perhaps switching dynamically depending on the circumstances, could adequately describe the theta phase code. However, we propose an additional scheme that could potentially reconcile these findings if they persisted after analyzing all relevant variables in the same dataset.

This third option would need to combine the spatial sweep’s constant place field sizes and phase precession slopes with the temporal sweep’s increase in theta trajectory length with running speed. The former property requires that sweeps are independent of running speed whereas the latter requires that they depend on it. While at first this appears to be a plain contradiction, there could be a solution if sweeps were made independent of the animal’s instantaneous running speed *v*(*t*), but dependent on their ‘characteristic running speed’ at each position 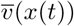. We refer to this as the behavior-dependent sweep model:

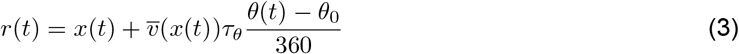

According to this model, at different phases of theta, the place cell population represents the positions that would have been or would be reached at certain time intervals in the past or future, like in the temporal sweep, but assuming the animal ran at the characteristic speed through each position, not at the instantaneous speed (Fig. 2A, B). Thus, the more stereotyped the behavior of the animal is, the more the behavior-dependent sweep model would share the temporal sweep model’s prediction that theta trajectory lengths increase proportionally with running speed. At the same time, the behavior-dependent sweep model is formally equivalent to the spatial sweep model (Eq. 1) with a theta distance that depends on the characteristic running speed at each position:

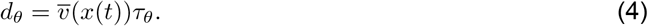

**Figure 2:**
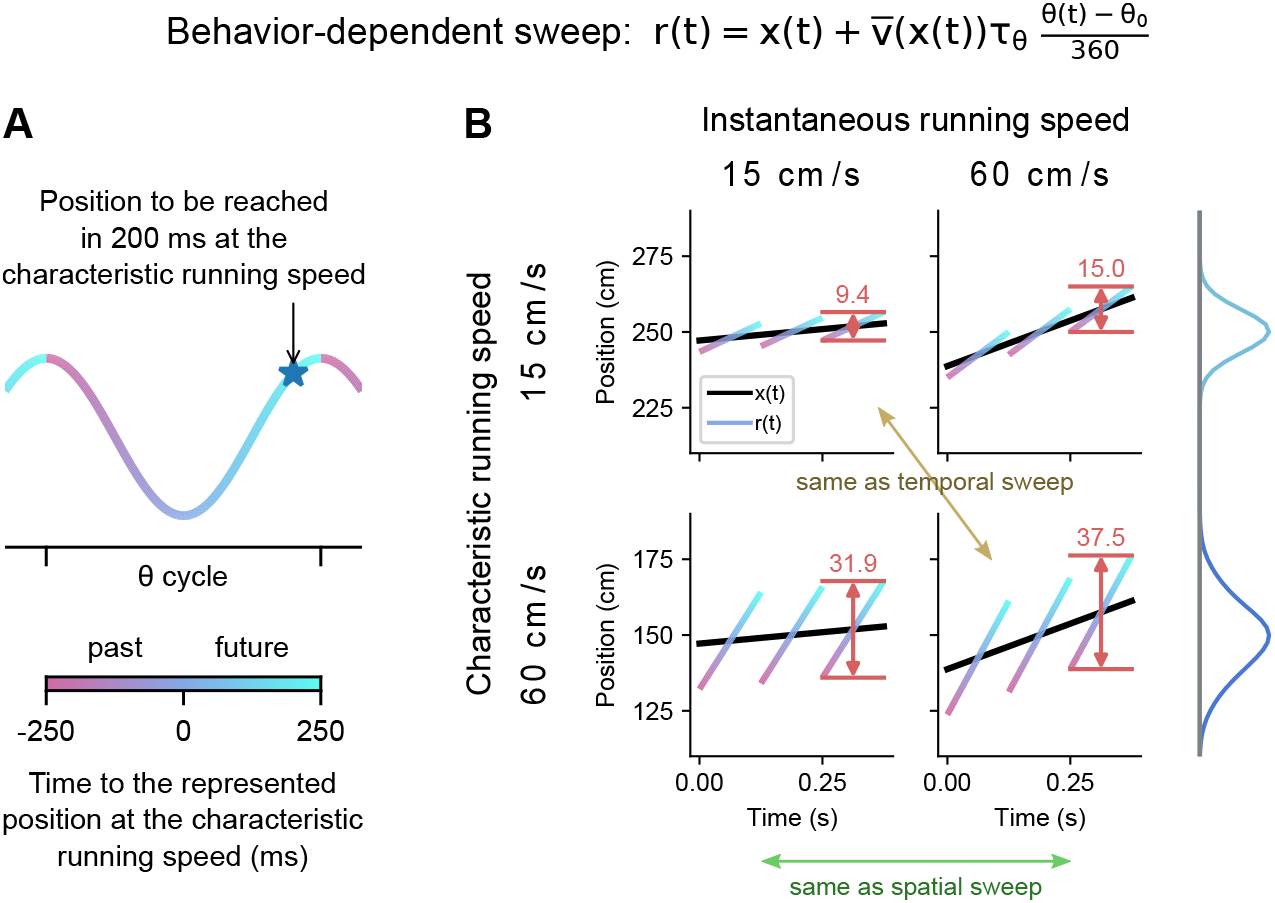
The behavior-dependent sweep integrates aspects of both spatial and temporal sweeps. Plotting convention as in Fig. 1A and C. **A**: Different phases of theta represent positions reached at different time intervals into the past or future assuming the animal ran at the characteristic speed at that location. **B**: Comparison of theta trajectory lengths in areas of low and fast characteristic running speed (rows) at low and fast instantaneous running speeds (columns). When both characteristic and instantaneous running speeds coincide, the behavior-dependent sweep model and the first order approximation of the temporal sweep model agree (ochre arrow). On the other hand, when changing the instantaneous speed at a given location, the behavior-dependent and spatial sweeps agree (green arrow). Place fields (right) are larger in areas of higher characteristic running speed.

Therefore, place fields, being fixed in space, would remain unaltered by the instantaneous running speed as in the spatial sweep model (Fig. 2B). However, place fields in areas of higher characteristic running speed would be larger and characterized by shallower phase precession slopes, something that has not been shown before to our knowledge.

In the following, we analyzed the effect of speed on theta trajectories and place field parameters in a single dataset to determine whether the apparently conflicting results can be observed simultaneously, and to gather data against which to quantitatively compare the model predictions of the three phase coding schemes.

### Theta trajectory lengths increase proportionally with running speed

To examine the effect of running speed on theta trajectory length, we re-analyzed publicly available simultaneous single unit recordings from CA1 of 6 rats running on linear tracks for water rewards at both ends. Three of the animals (Mizuseki et al., 2013) were running on a familiar track and the remaining three Grosmark et al. (2016), on a novel one. To only include periods with stable spatial representations and theta phase precession, we discarded the first five minutes of the recording on the novel track, after which time hippocampal place cells have developed stable place fields (Frank et al., 2004) and phase precession (Cheng and Frank, 2008).

We performed a Bayesian decoding of positions from the ensemble neuronal firing (Zhang et al., 1998; Davidson et al., 2009) within each theta cycle in overlapping 90°bins with a step size of 30°. Figure 3A shows examples of individual decoded theta trajectories belonging to periods of slow (below 20 cm/s) and fast (above 40 cm/s) running for one experimental session. The length of the trajectories was measured from best-fit lines. Despite the high level of variability, it was clear in the individual rats’ averages and in the grand average across animals that the length of trajectories increased roughly proportionally with running speed (Fig. 3B). We see the same effect if we average the decoded position probabilities over all cycles belonging to the same speed bin before fitting a line and calculating the theta trajectory length (Fig. 3C, D). For this analysis, we defined seven 20 cm/s wide overlapping speed bins starting at 2 cm/s in increments of 10 cm/s. Thus, our results on the dependence of theta trajectory lengths on running speed are consistent with those of Maurer et al. (2012) and Gupta et al. (2012), and support the temporal sweep model. Alternatively, the results could be accounted for by the behavior-dependent sweep model, if running behavior was sufficiently stereotyped across the track, a question to which we return below.

**Figure 3:**
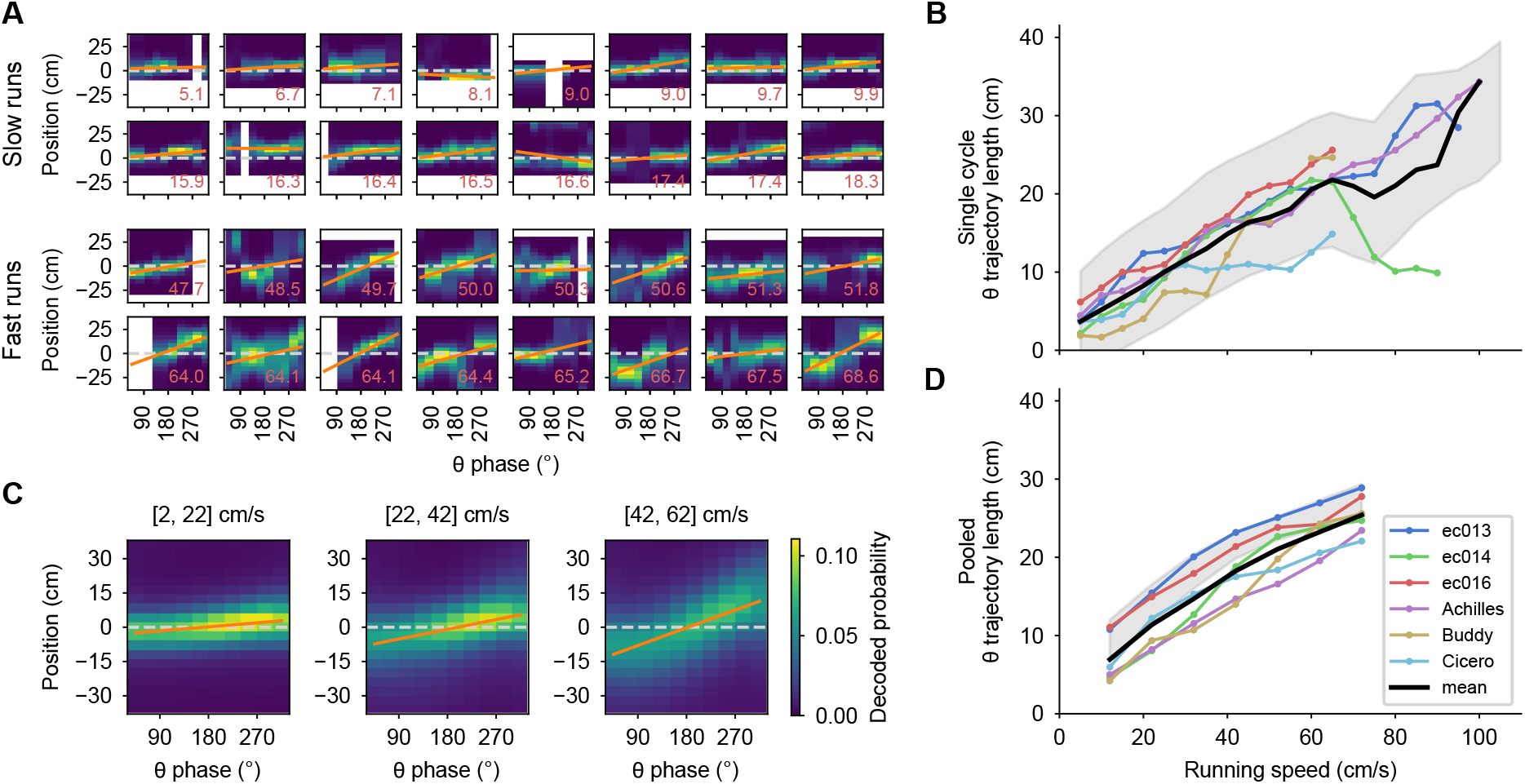
Theta trajectory lengths increase proportionally with running speed. **A**: A random sample of theta sequences from one experimental session, with position probabilities decoded from the population spikes. Zero corresponds to the actual position of the rat at the middle of the theta cycle; negative positions are behind and positive ones, ahead. Orange lines indicate linear fits, from which theta trajectory lengths are computed. White pixels correspond to positions outside of the track or phase bins with no spikes. The red number at the lower right corner indicates the running speed in cm/s. **B**: Moving averages for theta trajectory length for each animal using a 10 cm/s wide sliding window (colored lines). These averages are averaged again (thick black line) to obtain a grand average that weights all animals equally. Shaded region indicates a standard deviation around the mean of all of the underlying data points pooled across animals, which weight each theta sequence equally. This mean does not necessarily match the grand average since different animals contribute different numbers of theta sequences. **C**: Like A, but averaging the decoded probabilities across theta cycles belonging to the same speed bin, which is indicate above the panel. **D**: Like B for the averaged cycles, confirming the observation in B.

### Individual place field sizes remain constant with running speed

We next analyzed the effect of running speed on place field sizes. First, we calculated firing rates separately for periods with different running speeds using the same speed bins as before. The time spent by the animal in some spatial bins at some speeds was very low, leading to spurious firing rates of 0. Thus, we calculated place field sizes only where the field had been sufficiently well sampled (see Methods). Visual inspection of individual fields revealed that they tended to maintain their sizes at different running speeds (Fig. 4A). To quantify the relationship more systematically, we performed linear regression on place field size vs. speed for each field and examined the slopes of the best fitting lines (Fig. 4A, bottom row). Figure 4B shows that for each rat, slopes took on both negative and positive values, with no clear bias towards either (*p* > 0.3, Wilcoxon signed-rank test). Thus, we found no evidence for a systematic increase in place field size with running speed.

**Figure 4:**
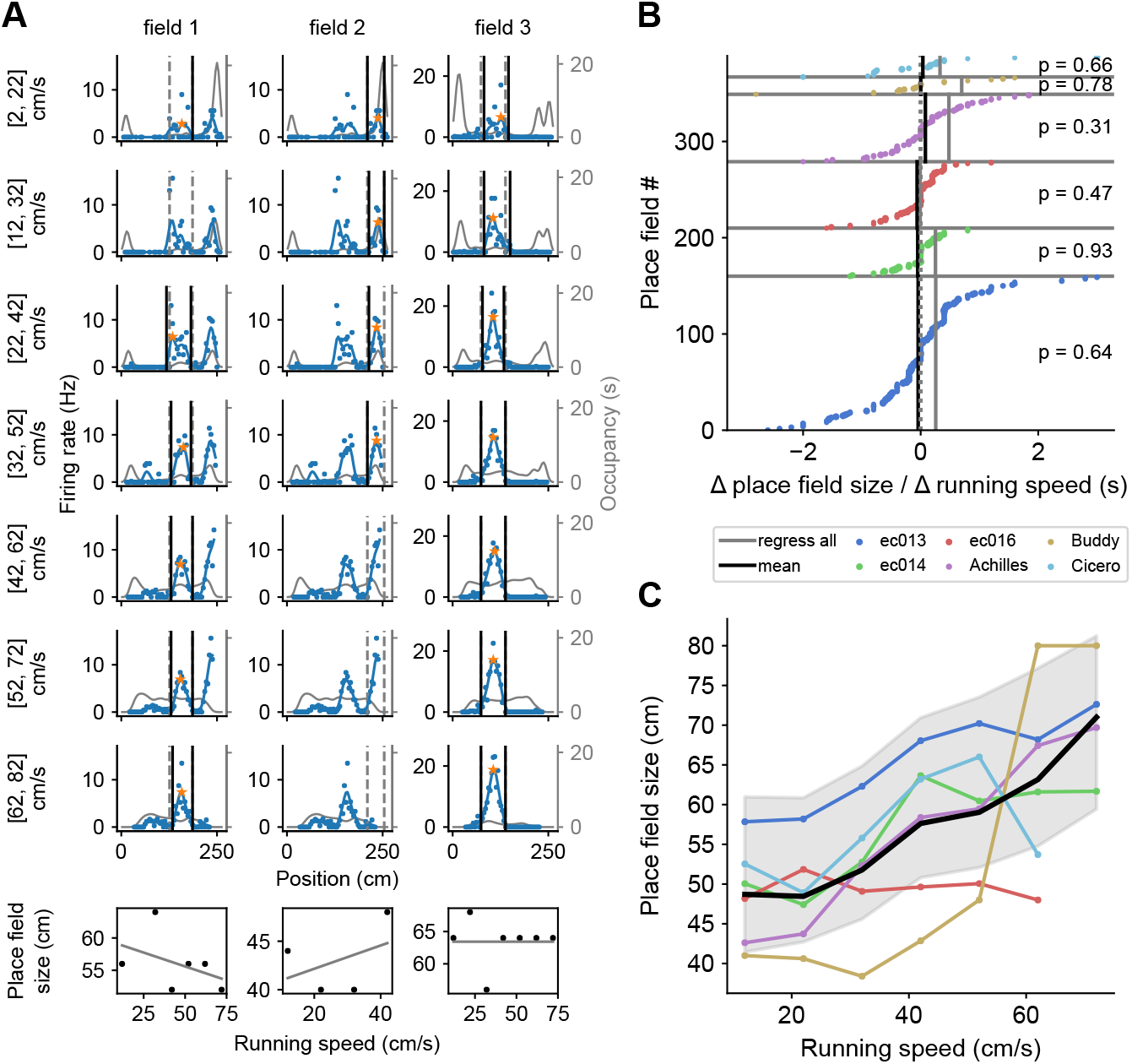
Place field size increases with running speed in pooled data, but not for individual fields. **A**: The size of a given place field remains roughly constant regardless of running speed. Examples from three individual place fields are shown (one per column). Dashed gray lines represent the extent of the place fields calculated from the complete set of all spikes at all running speeds. Black lines mark the extent of the place field calculated for each speed bin. Where only one end of the place field could be determined (e.g., field 2, third and fourth rows), place field size was calculated as twice the distance from the field’s peak (orange star) to the detected field end (see Methods). Thin gray lines represent the occupancy (time spent) per spatial bin (axis on the right). At the bottom, linear regressions on place field size vs. running speed for each place field. **B**: The slopes from such linear regression of place field size vs. running speed for all place fields, sorted for each animal. The slopes are not significantly different from zero (indicated p values). The size of the dot reflects the number of data points that contributed to the regression. The black vertical lines indicate the weighted averages of these slopes for each animal. The gray lines indicate the slope of the regression calculated by first pooling together data points from all place fields for each animal. **C**: Remarkably, when pooling place field sizes across fields, the average field size generally increases as a function of running speed. Colored lines represent individual animals, and the thick black line averages over them. Shading represents a standard deviation around the mean of all data points pooled across animals.

Intriguingly, however, when place field sizes were pooled across fields, they tended to be larger at higher speeds as shown by the mean place field size in each speed bin (Fig. 4C) and statistical analysis (*p* < 0.03 for place field sizes pooled across fields increasing with speed for 5 of the 6 animals, no significant relationship for the remaining one; Wald Test with t-distribution of the test statistic). This increase in pooled place filed sizes is consistent with the temporal sweep model, whereas the lack of within-field changes supports spatial sweeps. Both findings, however, appear consistent with behavior-dependent sweeps.

We also re-ran the earlier theta trajectory length analysis (Fig. 3D) restricted to the areas and speeds included in the place field analysis without observing substantial changes (*SI Appendix*, Fig. S2).

### Phase precession slopes of individual place fields remain constant with running speed

Similar results were obtained when analyzing phase precession. Since the temporal sweep model predicts very steep phase precession clouds for low speeds (e.g., Figure 1D), we devised a method that could capture such steep phase precession slopes using the orthogonal regression (see Methods). First, we calculated phase precession slopes separately for each running speed bin by pooling spikes based on the instantaneous running speed of the animal when the spikes were emitted. Figure 5A shows the results of this analysis for three example place fields. An analysis of all place fields (Fig. 5B) revealed that phase precession slopes of individual fields did not change systematically with running speed for any of the six animals (*p* > 0.05, Wilcoxon signed-rank test). Remarkably, however, when phase precession slopes were pooled across fields in each speed bin, the slopes at slower speeds tended to be lower than at faster speeds as reflected in the mean phase precession slope at each speed (Fig. 5C) and statistical analysis (*p* < 0.002 for phase precession slopes pooled across fields increasing with speed for 5 of the 6 animals, no significant relationship for the remaining one; Wald Test with t-distribution of the test statistic).

**Figure 5:**
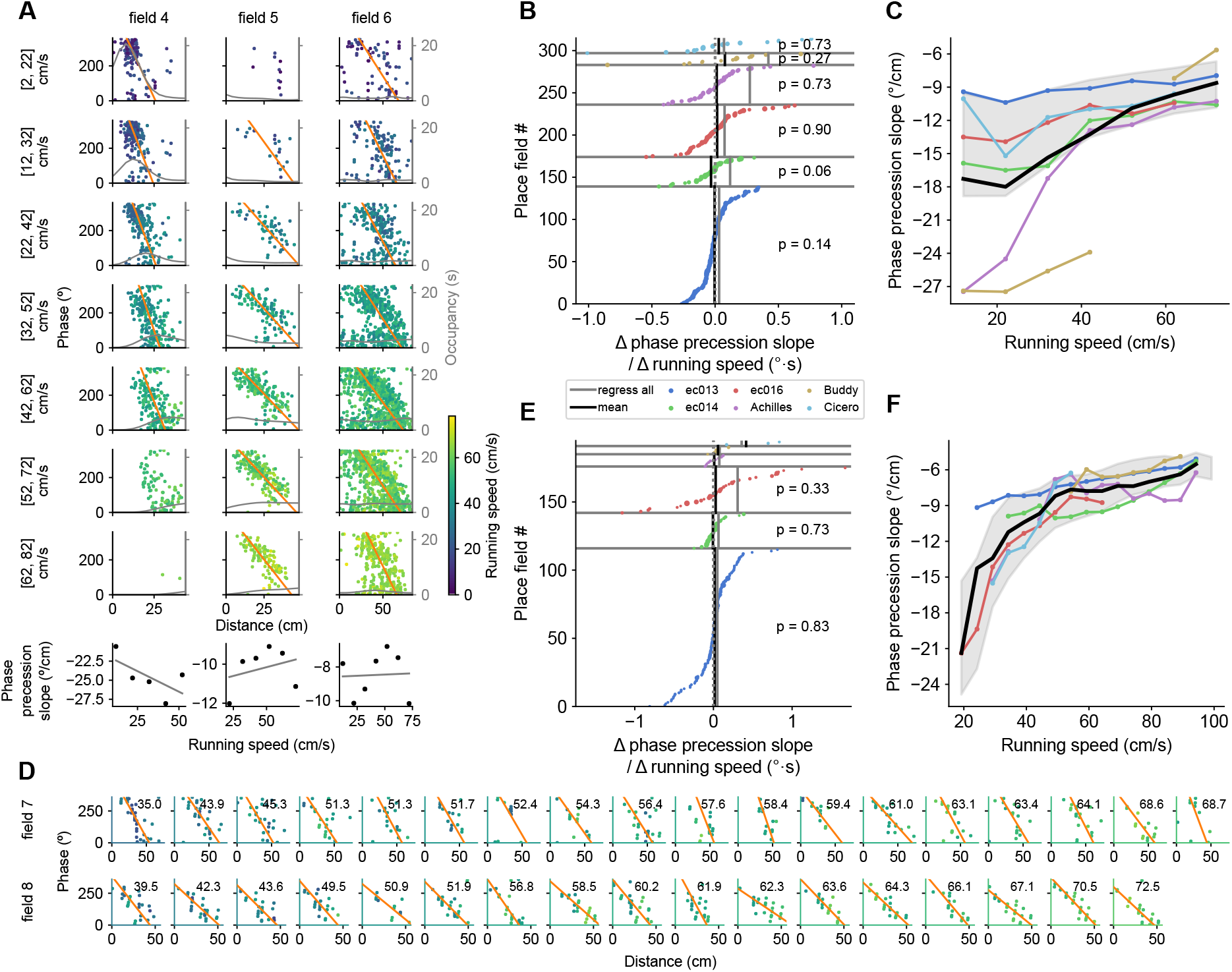
Phase precession slope increases with running speed in pooled data, but not for individual fields. **A**: Example phase precession slopes at different speeds for three fields. Instantaneous speeds when the spikes where emitted are color coded. Thin gray line displays the occupancy. At the bottom, linear regressions on phase precession slope vs. running speed for each place field. **B**: As in Fig. 4B for the slopes of the linear regressions on phase precession slope vs. running speed for individual fields. **C**: As in Fig. 4C for phase precession slopes pooled across fields for each speed bin. **D**: Example phase precessions in single passes through two place fields (rows) sorted by running speed (upper right corner, in cm/s). Color code as in A. **E, F**: Same as in B and C but for single-trial phase precession slopes. The means for each animal in F were calculated as a moving average on a 10 cm/s wide sliding window.

In the analysis just described, when rats typically accelerate or decelerate through some field, different parts of the field’s phase precession cloud are sampled at different running speeds. This can be seen in the first field (no. 4) in Fig. 5A. Since the rat was accelerating through the field, spikes emitted at lower and higher speeds appeared more frequently at the beginning and end of the field, respectively. Phase precession clouds are known to have some curvature (Souza and Tort, 2017), and so sampling different parts of the field when accelerating or decelerating could lead to a spurious relationship between speed and phase precession slope.

To avoid this effect, we decided to further analyze phase precession slopes in single passes through a field (Schmidt et al., 2009). We only included passes that occurred at approximately constant speed, i.e., passes with a coefficient of variation in speed lower than 0.3. Fig. 5D shows examples of individual passes through two fields, sorted by running speed. Note how, notwithstanding variability in the slopes, there is no systematic relationship with running speed, even though the speeds nearly double. There was no significant relationship between phase precession slope and speed for individual fields in any of the three animals with sufficient data for this analysis, i.e., more than 10 fields (Fig. 5E). Yet again, when we pooled phase precession slopes across place fields, we saw a systematic increase in phase precession slopes with speed in all animals (*p* < 0.05, Wald Test with t-distribution of the test statistic; Fig. 5F). This hyperbolic increase in pooled phase precession slopes supports the temporal sweep model, whereas the absence of within-field increases in phase precession slopes with running speed is consistent with the spatial sweep model. Both experimental findings, however, appear consistent with behavior-dependent sweeps.

### Theta trajectories and place field parameters vary across the track based on characteristic running speed

We found that theta trajectory lengths, place field sizes and phase precession slopes increase with running speed, if data from different positions along the track or place fields are pooled in the analysis (Fig. 3B, D; Fig. 4C and Fig. 5C, F), but not when fields are analyzed individually (Fig. 4B and Fig. 5B, E). While these simultaneous findings might seem counter-intuitive, the behavior-dependent sweep model offers a simple explanation: Short theta trajectories and small place field sizes could occur where animals tend to run slow, contributing to the low speed averages, and long theta trajectories and large place field sizes could occur where animals tend to run fast, contributing to the fast speed averages (Fig. 6A).

**Figure 6:**
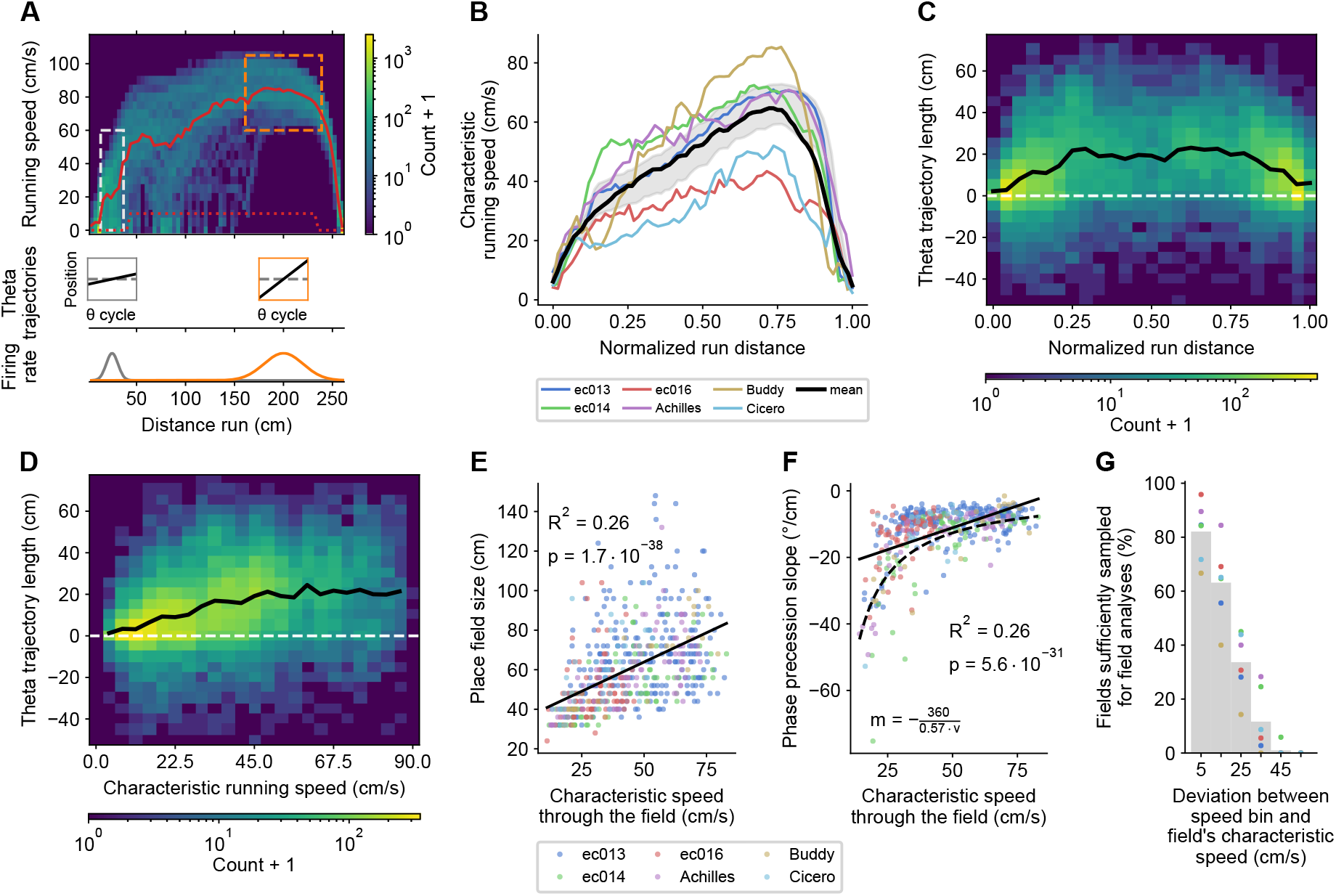
Structured place field and theta trajectory heterogeneity correlates with characteristic speed. **A**: Potential explanation for the increase in average theta trajectory length, place field size and phase precession slopes with speed despite the lack of systematic within-field changes. The histogram shows the distribution of running speeds by positions for rightward runs in one experimental session. The thick red line is the characteristic speed, defined as the mean speed after discarding running speeds below 10 cm/s in the center of the track (exclusion criteria indicated by dotted red line). Running speeds tend to cluster around the characteristic speed. We hypothesize that short theta trajectories and small place fields occur in positions with low running speeds, while long theta trajectories and large place fields occur where animals run faster. **B**: Average characteristic speed as a function of the normalized distance from the start of the run for each animal (colored lines) and grand average across animals (thick black line). Shaded region as in Figure 3B. **C**: Histogram of individual theta trajectory lengths across the track for all animals combined and their mean (black line). Negative values correspond to theta trajectories going opposite to the running direction. **D**: Histogram of theta trajectory lengths vs. characteristic running speed at the position where trajectories were observed. **E**: Place fields are larger and **F**: have shallower phase precession slopes, the higher the mean characteristic speed through the field is. **G**: The proportion of fields sufficiently sampled for place field analysis at a certain speed bin falls steeply with the deviation between the speed bin and the mean characteristic speed through the field. Grey bars indicate averages across animals.

Specifically, this explanation predicts that 1. running speeds differ systematically along the track, 2. theta trajectory lengths and place field parameters also differ systematically along the track, being proportional to the characteristic speed at each position and, 3. different place fields contribute differentially to low and high speed averages.

To test prediction 1, we calculated the characteristic speed for each spatial bin along the track. This was defined as the mean speed after discarding running speeds below 10 cm/s in the center of the track, to prevent atypical pauses in running from disproportionately distorting the mean values. The upper panel in Fig. 6A shows the characteristic speed for rightward runs in one sample experimental session and Fig. 6B shows averages across running directions and sessions. All animals showed a similar asymmetric inverted U-shape relationship between characteristic speed and position along the track.

In support of prediction 2, we also observed a striking inverted U-shape relationship between individual theta trajectory lengths and position along the track (Fig. 6C), suggesting a link to characteristic running speed. Indeed, theta trajectory length increased roughly proportionally with characteristic speed for a large range of speeds (Fig. 6D). Furthermore, we found that place field sizes and phase precession slopes also displayed inverted U-shape relationships to position along the track and increased significantly with the characteristic speed through the field when pooling fields from all animals (Fig. 6E, F; *p* < 10^−30^, Wald Test with t-distribution of the test statistic) and for most animals individually (*SI Appendix*, Fig. S3).

Finally, we confirmed prediction 3 that place fields mostly contributed data points to the place field or phase precession slopes averages in speed bins near the characteristic speed through the field. Only 12% of fields were sufficiently sampled to contribute points to the analyses at speeds differing by 35 cm/s from the fields’ characteristic speeds, falling to 0% at speed differences larger than 55 cm/s (Fig. 6G).

Taken together, these findings reconcile the speed-dependence of average theta trajectory lengths, place field sizes and phase precession slopes with the speed-independence of individual fields, and confirm the predictions of the behavior-dependent sweep model.

### The behavior-dependent sweep model accounts best for all experimental results

Our preliminary analyses suggest that neither the spatial nor the temporal sweep models can account for all of the experimental observations, whereas the behavior-dependent sweep appears consistent with all of them. However, these analyses might involve too many simplifying assumptions about how running speeds, theta coupling and place field parameters are co-distributed. To address this issue, we developed phenomenological models of phase coding that use the experimental LFP and tracking data, but generate spikes based on the different theta coding schemes. The outputs of these models were then analyzed in the same way as the experimental data (Fig. 7). This further allowed us to examine whether any of the models could, using a single set of parameters, provide a unified account of the observed phenomena.

**Figure 7:**
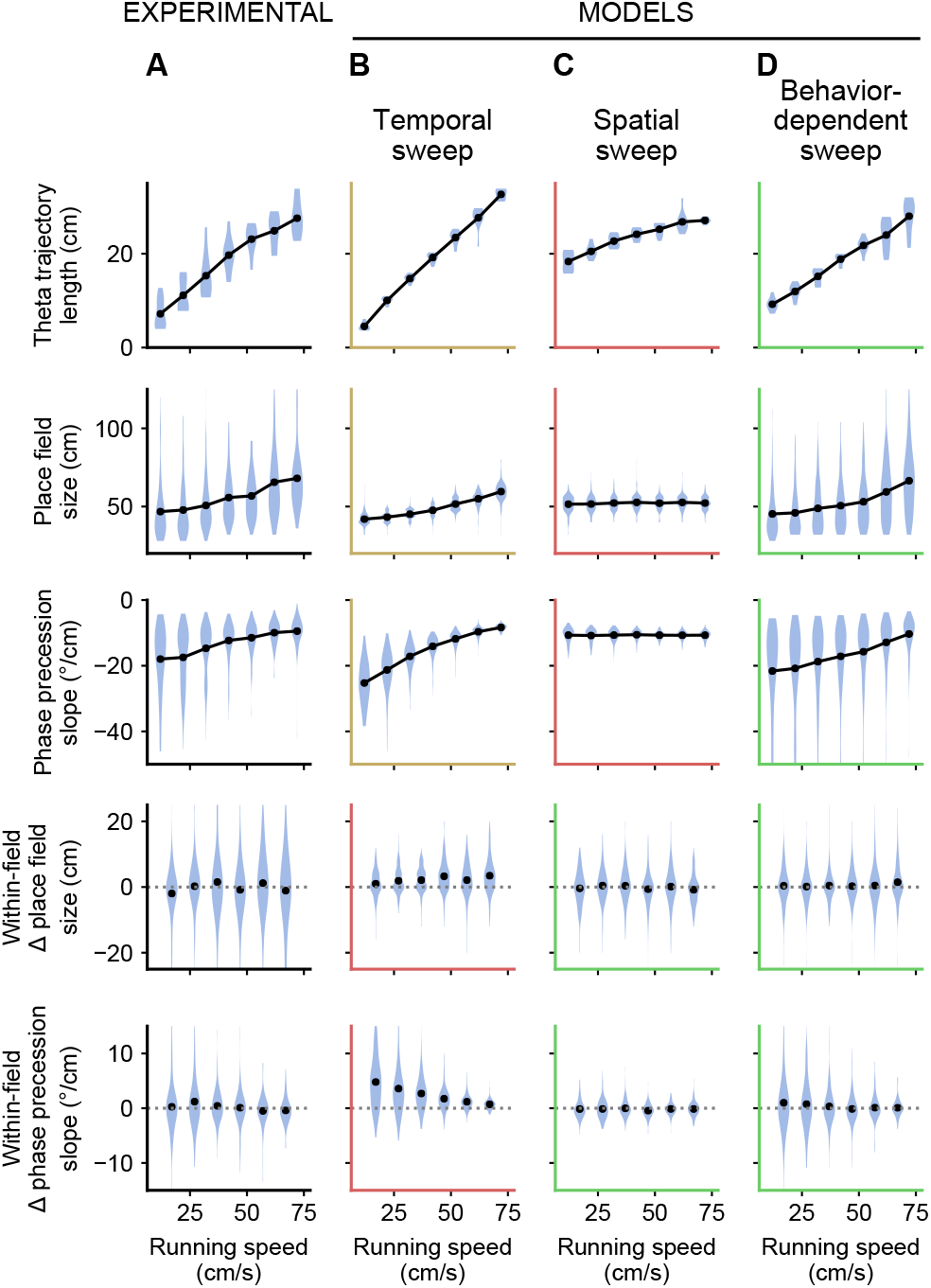
Only the behavior-dependent sweep model accounts for all experimental observations. **A**: Summary of experimental results from all animals. Only one animal had more than three sessions. For this animal, we chose three sessions at random so that it would not dominate the results. From top to bottom row: theta trajectory lengths, place field sizes and phase precession slopes pooled according to running speed, and the change in place field size and phase precession slope from one speed bin to the next for each field. Black dots represent mean values. **B**: The temporal sweep model provides a qualitative fit (ochre axes) to theta trajectory lengths, place field sizes and phase precession slopes across the population, but incorrectly predicts (red axes) increases in individual place field sizes and phase precession slopes. **C**: The spatial sweep model accounts for (green axes) the lack of within-field increases in place field sizes and phase precession slopes. However, the increase in theta trajectory length with running speed is not large enough and place field sizes and phase precession slopes remain flat with running speed. **D**: The behavior-dependent sweep model captures both the population average and within-field effects, providing a good agreement with all experimental results.

The temporal sweep model could be fitted to roughly capture the observed increase in theta trajectory lengths, place field sizes and phase precession slopes pooled by running speed, but it incorrectly predicted changes in individual fields with speed (Fig. 7B). By contrast, the spatial sweep model could account for the speed-independence of individual fields, but severely undershot the increase in theta trajectory length with running speed and failed to capture the increases in place field size and phase precession slopes (Fig. 7C).

For the behavior-dependent sweep model, *τ_θ_* is the crucial parameter determining theta trajectory length and place field measures. We estimated this parameter from the experimentally observed phase precession slopes, which display the most straightforward relationship to it. In particular, the model predicts an inverse relationship between phase precession slope *m* and characteristic running speed 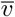 (Methods). Assuming that the theta phase precesses over 360°, 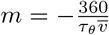. Our data seems to conform to this relationship (Fig. 6F). Indeed, when the inverse of the slope is plotted against the characteristic speed, they are roughly proportional to each other up to 75 cm/s (*SI Appendix*, Fig. S3C). Fitting this data with the derived function, we obtained an estimate of *τ_θ_* = 0.57 s.

Remarkably, with this value, the behavior-dependent sweep model provides a good fit to the rest of experimental results. The model reproduces the lack of within-field changes with running speed (Fig. 7D), since the extent of the spatial sweeps through each portion of the track is fixed. The model also captures the experimentally observed increase in theta trajectory lengths, place field sizes and phase precession slopes with running speed in data pooled by running speeds.

A further, more subtle, prediction of this model is that fields will skew differently depending on whether the animal is on average accelerating or decelerating through the field. We therefore analyzed the relationship between mean acceleration and place field skew in the experimental data and found a good qualitative fit with the predictions from the model (*SI Appendix*, Fig. S4A). Negatively skewed fields Mehta et al. (2000) were observed at all acceleration values, but positively skewed fields occurred mainly at positive accelerations, resulting overall in a significant correlation between mean acceleration and skew (p < 10^−4^; Wald Test with t-distribution of the test statistic). A similar observation was made by Diba and Buzsáki (2008), who reported positive skews at the beginning of the track, where rats are usually accelerating, and negative skews towards the end, where rats decelerate.

In summary, we find that the behavior-dependent sweep model provides the best fit to our experimental data, and does so in a realistic setting, using the experimental LFP and tracking data, and with a single set of model parameters.

## Discussion

While research over the past four decades has shed light on numerous aspects of the hippocampal code, the exact relationship between represented position, theta phase, and the animal’s physical location has remained ambiguous. The expression of this relationship in theta sequence trajectories at the population level and in theta phase precession at the single-cell level are two sides of the same coin, but the two levels have seldom been explored side by side. Here, we illustrated that established findings in the field at these different levels are in apparent contradiction with one another (Maurer et al., 2012; Gupta et al., 2012; Geisler et al., 2007; Diba and Buzsáki, 2008; Schmidt et al., 2009; Huxter et al., 2003). We reconciled these findings by demonstrating that even though individual place fields do not change with instantaneous running speed, their size and phase precession slopes vary according to the animal’s characteristic running speed through these place fields. In particular, regions of slow running are tiled with smaller place fields with steeper phase precession, corresponding to shorter theta trajectories at the population level, whereas regions of fast running are covered by larger place fields with shallower phase precession, corresponding to longer theta trajectories. Based on these observations, we proposed a novel theta phase coding scheme, the behavior-dependent sweep. In this model, the spatial extent of the theta oscillation sweep varies across the environment in proportion to the characteristic running speed at each location. As a result, the sweeps can be seen as going through the positions that were or will be reached at certain time intervals in the past or future, assuming the animal runs at the speed characteristic of each place, as opposed to at its actual instantaneous speed. This coding scheme thus integrates aspects of both the spatial and temporal sweeps.

### Relationship to previous studies on the speed-dependence of the hippocampal code

Why were the relationships between characteristic speed, hippocampal sweeps and place field parameters missed by previous studies? Two previous studies (Maurer et al., 2012; Gupta et al., 2012) pooled theta trajectories based exclusively on speed, therefore masking the role of position in mediating the relationship between speed and theta trajectory length. In other studies, the dependence of place field parameters on the characteristic running speed through the field was missed because analyses focused on relative measures between pairs of cells (Geisler et al., 2007; Diba and Buzsáki, 2008) or individual fields (Schmidt et al., 2009; Huxter et al., 2003; Maurer et al., 2012) that remain unaffected.

We note that one other study (Chadwick et al., 2015) previously proposed a model to account for the increase in theta trajectory length with running speed despite stable phase-precession. The model achieved this by increasing the precision of the theta phase versus position relationships of cells with running speed. This feature, however, results in unusually noisy phase-precession at low speeds (*SI Appendix*, Fig. S5 and see Appendix for results on a related model), which we did not observe in the data. Furthermore, this model cannot account for the dependence of the average place field sizes and phase-precession slopes on characteristic running speed.

On the other hand, two models which are consistent with this increase in pooled place field parameters did not address theta phase coding. Slow feature analysis attempts to extract slowly varying signals from the sensory input, as these signals are deemed to be more useful, and has been found to account for different properties of place, head direction and view cells (Franzius et al., 2007). Extracting place-specific features that vary on the order of several seconds, for instance, would lead to larger place fields at locations of higher speeds, as we have observed. This observation is also consistent with the successor representation model of place-fields (Stachenfeld et al., 2017). In this model, the activity of a cell is determined by the expected discounted future occupancy of the cell’s preferred location, which proves useful for planning. Thus, a cell with a preferred location where average running speed is higher would begin firing relatively further away from that place. The behavior-dependent theta sweep model shares this prediction with these two models, and further accounts for theta-phase and trajectory relationships.

### Behavior-dependent sweeps are advantageous for prediction and planning

The temporal compression of trajectories within theta cycles has been suggested to support a hippocampal role in prediction and planning (Lisman and Redish, 2009; Erdem and Hasselmo, 2012; Bush et al., 2015; Vanderwolf, 1969), particularly at the later phases of theta (Hasselmo et al., 2002). We can thus compare our three coding schemes in terms of their benefit for prospection and decision making. Representing future positions based on the distance to them, as in the spatial sweep, could allow an animal to calculate the speed necessary to reach those positions in a given amount of time. This representation might also be convenient because our bodies and sensory organs limit the spatial range at which we can interact with the world. In contrast, looking forward based on an interval of time rather than space, as in the temporal sweep model, could increase the code’s efficiency by matching the extent of the look-ahead to the speed of progression through the environment. For instance, while driving a car on a highway, we might have to look ahead by several hundred meters, since those positions are just seconds away, while such a distance is not useful when strolling in a park. Plans made too far in advance are very likely to be discarded upon unexpected changes or as new information comes to light, which is conceivably related to the use of a discount factor in reinforcement learning models (Sutton and Barto, 1998). Temporal sweeps also seem more plausible for phase-coding in non-spatial domains (Pastalkova et al., 2008; Lenck-Santini et al., 2008; Robinson et al., 2017; Terada et al., 2017; Aronov et al., 2017). However, temporal sweeps require a fine control by the instantaneous running speed, and there are indications that the hippocampal network lacks the the flexibility required to modify the time lags between pairs of cells at the theta timescale (Diba and Buzsáki, 2008; Shimbo et al., 2021).

The behavior-dependent sweep might thus represent the best solution compatible with network constraints by taking advantage of the relative simplicity of spatial sweeps while still enjoying the benefits of temporal sweeps in situations in which behavior is stereotyped. Thus, the look ahead on a highway remains a few hundred meters even if on occasion we are stuck in a traffic jam. Finally, behavior-dependent sweeps could make predictions of future positions more robust through the averaging of past experience.

Despite the potential benefits of behavior-dependent sweeps mentioned here, these sweeps are probably best construed as a default or baseline mode of operation, which could be further modulated by ongoing cognitive demands and task contingencies (Wikenheiser and Redish, 2015; Shimbo et al., 2021; Zheng et al., 2020).

### Look-ahead times can help explain place field sizes

The link between place fields and prospective representations at the theta timescale allows us to set a lower bound on what place field sizes would be useful for a given task. For sweeps that are centered around the current position, a *τ_θ_* of ~ 0.6 s would mean that theta sweeps on average end in positions arrived at in ~ 300 ms. Representing those future positions would be of use only if there was enough time for the animal to modify its behavior in response to them. Reaction times for rats performing simple and choice tasks range between 150 ms and 500 ms (Baunez et al., 2001; Brown and Robbins, 1991), and stop signal reaction times stand at around 360 ms (Eagle et al., 2009). Therefore, place field sizes associated with a look-ahead time of 300 ms probably stand towards the lower end of what can be used to control behavior through rapid decision-making. However, place fields (Kjelstrup et al., 2008) increase in size along the dorsoventral axis of the hippocampus. The largest place fields recorded in rats stand at about 10 m, which would correspond to representing a time window of > 10 s. Thus, we can conceive of a hierarchy of subregions corresponding to predictions at different temporal scales.

### Potential mechanisms of behavior-dependent sweeps

The dependence of place field parameters and theta trajectories on characteristic speed raises the question of how and when this speed might be computed in the brain, and how it is used to control hippocampal sweeps. One intriguing possibility is that the animal can estimate the speed at which it will typically traverse some region of an environment based on general knowledge of similar environments (e.g., knowing that speed will tend to be lower around boundaries). Another possibility is that the characteristic speed is taken to be the speed of the firsts traversals through the environment, in which theta sequences are reportedly lacking or reduced (Feng et al., 2015; Tang et al., 2021), and then updated incrementally.

The latter suggestion fits naturally with network connectivity models of phase precession (Jensen and Lisman, 1996; Tsodyks et al., 1996; Drieu and Zugaro, 2019).In these models, theta sequences are produced by the propagation of activity through the population, perhaps owing to the recurrent connectivity in area CA3 (Cheng, 2013; Azizi et al., 2013). The network is cued with the current position and recalls upcoming positions. As the animal moves forward, positions will activate at earlier phases of the theta cycle, producing phase precession. In this family of models, it is easy to imagine sweeps varying based on average speed. Place cells could be connected with a higher synaptic strength the closer in time they become activated, leading to stronger and further reaching connectivity in areas through which the animals run faster on average. If the speed of propagation in the network is then made dependent on synaptic strength, activity would propagate further in the network within each theta cycle where speed had been higher, producing longer theta trajectories in those areas.

### Predictions and outlook

In the dataset analyzed, changes in running speed arose naturally from animals running on a finite track, but were not controlled by the experimenter. Further confirmation for the behavior-dependent sweep model could come from experiments in more complex environments that can produce variable running speeds at different specific locations (Kropff et al., 2021), such as slowing down in previously fast segments and speeding up in slow ones. In particular, it would be interesting to dissociate characteristic running speed and distance from environmental boundaries, since boundary vector cell (Barry et al., 2006) and Laplace transform (Sheehan et al., 2021) models predict that distance to the border or trial start, rather than characteristic running speed, are what determines place field size.

Although we have focused primarily on the effect of speed, it is reasonable to also expect acceleration to have an effect on theta phase coding. We discuss three specific mechanisms. First, acceleration could introduce a second-order term in the equation for *r*(*t*) such that theta trajectories curve in areas through which the animal typically accelerates or decelerates. This might cause trajectories in areas of positive acceleration to reach further ahead than behind, and trajectories in areas of negative acceleration to reach further behind than ahead, consistent with the findings of Gupta et al. (2012) and Bieri et al. (2014). Second, acceleration leads to different characteristic running speeds in different parts of a place field. Our model predicts that this would cause place fields to skew differently in areas of positive and negative acceleration, a prediction for which we have found some preliminary experimental support (*SI Appendix*, Fig. S4A). Additionally, the model indicates that changes in characteristic speeds through fields cause corresponding changes in theta phase precession. If characteristic speed increases across a field, the slope of phase precession would become increasingly shallow, resulting in concave phase precession clouds, whereas if characteristic speed decreases across the field, the slope of phase precession would be increasingly steep, producing convex “banana shaped” phase precession clouds (Souza and Tort, 2017; Yamaguchi et al., 2002). Future studies are required to test these two predictions, since the range of accelerations in the current dataset was relatively small, specially for positive values, and strong acceleration and deceleration were concentrated over small regions at the start and end of the track, respectively. Third, theta frequency has recently been shown to increase with positive acceleration, but not deceleration or speed (Kropff et al., 2021). One might wonder whether shorter theta cycles due to positive acceleration, rather than the lower running speed, could explain the shorter theta trajectories we observed at the beginning of the track. However, we also observe shorter theta trajectories at the end of the track, where running speeds are also lower, but acceleration is negative and theta cylces are longest (Fig. 6B, C). Furthermore, as shown by (Kropff et al., 2021), the increase in theta frequency is accompanied by an increase in the intrinsic firing frequency of cells which would provide a compensatory effect on theta trajectories. Even without such compensation, the reduction in theta trajectory length due to shortened theta cycle is small and insufficient to account for the observed effect.

Lastly, our behavior-dependent sweep model could be applied to describing the activity of place cells in two-dimensional environments, and potentially also that of grid cells, since theta phase coding has been demonstrated in both of these cases (Huxter et al., 2008; Jeewajee et al., 2014; Hafting et al., 2008). For grid cells, our model predicts that sizes of individual firing fields within the grid will vary according to running speeds characteristic of the corresponding maze regions. By contrast, the distance between firing field peaks would remain constant, since the locations of the underlying true fields remain fixed in the model.

In conclusion, our analyses have highlighted the heretofore unresolved tension lurking in the field between experimental results and interpretations explicitly or implicitly supporting spatial and temporal sweeps. Our novel interpretation reconciles apparently contradictory findings by making spatial sweeps dependent on the average behavior at each position, combining features and benefits of both kinds of sweep. The model advances our understanding of how the mammalian brain quantitatively represents past, present and future spatial positions for learning and planning.

## Methods

### Mathematical description of coding schemes

#### Spatial sweep

In the spatial sweep model, the represented position at time *t*, *r*(*t*) is given by

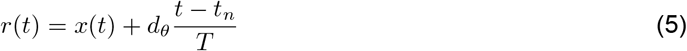

where *x*(*t*) indicates the actual position of the animal, *d_θ_* represents the extent of the spatial sweep, *T* is the period of the theta oscillation and *t_n_* is the time for which *r*(*t*) = *x*(*t*) within the current theta cycle.

In order to derive an expression for the length of theta trajectories, we can perform a Taylor series expansion around *t_n_*:

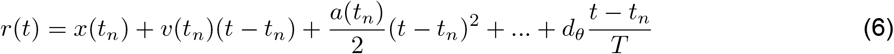

where *v* and *a* are the speed and acceleration respectively. Since (*t − t_n_*) is small, we can ignore the effects of the acceleration and higher order derivatives and approximate the represented position as:

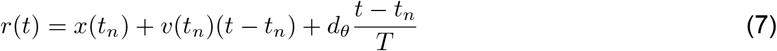

Here, the first two terms reflect the actual movement of the animal and the last one represents the sweep across positions starting behind the current position of the animal and extending ahead.

From here we can calculate the length of theta trajectories, which encompasses both the change in the actual position of the animal due to its movement during a theta cycle and the sweep:

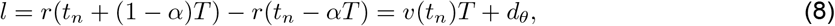

where *α* is the relative position within the theta cycle for which *r*(*t*) = *x*(*t*). The length of theta trajectories has an offset of *d_θ_* and is only weakly dependent on speed since the speed term is multiplied by the relatively small theta period.

Lastly, we can easily provide a mathematical description of the phase precession slope by noting that the phase will precess over *d_θ_*. Assuming a phase precession range of 360° we obtain the constant:

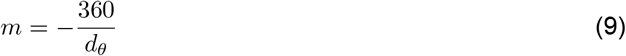

#### Temporal sweep

For the temporal sweep model, the represented position is given by,

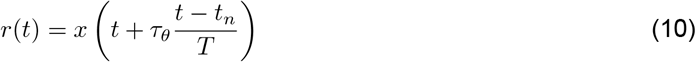

where *τ_θ_* is the extent of the temporal sweep. Again, we can perform a Taylor series expansion around *t_n_*:

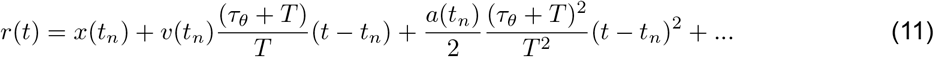

And discarding the effects of higher order derivatives:

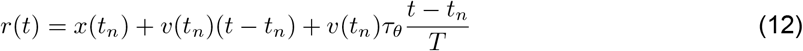

where again the first two terms reflect the movement of the animal and the last one, the sweep. In this case, the length of the theta trajectories is directly proportional to running speed:

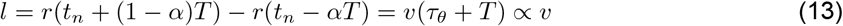

From the first order approximation of *r*(*t*) in Eq. 12 we see that the spatial extent of the sweep is *v*(*t_n_*)*τ_θ_*, that is, the space that will be covered at the current speed over the temporal window defined by *τ_θ_*. Thus, the phase precession slope is:

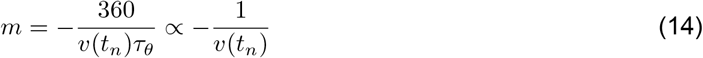

which is inversely proportional to running speed, leading to an increasingly steep phase precession as speed goes to 0.

#### Behavior-dependent sweep

A way of integrating aspects of spatial and temporal sweeps is to make the extent of spatial sweeps directly proportional to the average running speed at each location. Thus, we set 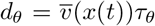 in Eq. 5:

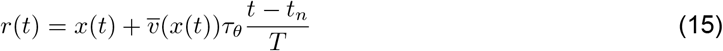

Note that second term representing the sweep is equivalent to the sweep term in Eq. 12, but using the average speed through the current location as opposed to the actual instantaneous speed at which the animal is running.

In order to account for the variability in the data, we consider the case where each cell operates based on a noisy measure of *d_θ_*. Thus, each cell represents a slightly different position, *r_i_*(*t*):

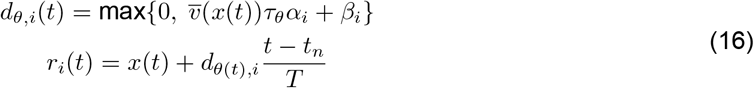

with *α_i_* = *N* (1, *μ_mul_*) and *β_i_* = *N* (0, *μ_add_*), fixed for each cell.

Proceeding as above, we can derive equations for the theta trajectory length

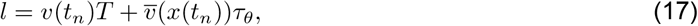

which is roughly proportional to average running speed, and phase precession slope

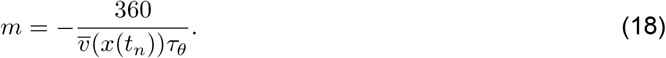

### Data analysis

We re-analyze data from two publicly available data sets: Mizuseki et al. (2013) and Grosmark et al. (2016), described in Mizuseki et al. (2009, 2014) and Grosmark and Buzsaki (2016) respectively. Briefly, 7 male Long-Evans rats ran on linear tracks of various lengths for water rewards at both ends. Three rats were running on a familiar track and 4 on a novel track. One of the rats running on a novel track was excluded from the study due to issues with local field potential recordings (see below). The first 5 min of the recordings in a novel track were discarded, since we wanted to only include periods with stable spatial representations, and place fields of hippocampal place cells might be unstable in the first 5 min of novel experience (Frank et al., 2004).

Neural activity was recorded using multi-shank silicon probes targeting region CA1 of dorsal hippocampus. After spike sorting, cells were classified as principal neurons or interneurons based on monosynaptic interactions, wave-shapes and burstiness.

#### Local field potential

We filtered the local field potential (LFP) with a Butterworth filter of 3rd order between 4 and 12 Hz. We then estimated theta phase from the signal’s Hilbert transform and shifted it by 180° so that 0° and 360° roughly correspond to the peaks of the filtered LFP. Individual theta cycles were then defined as spanning the range from 0° to 360°. Periods of significant theta oscillation were identified as those with an instantaneous amplitude above the 97th percentile of amplitudes in a shuffled surrogate signal. For creating this surrogate, we high-pass filtered the LFP at 1 Hz with another Butterworth filter of 3rd order and randomly permuted the time stamps. All of the analyses carried out throughout the paper deal exclusively with periods of significant theta oscillation.

In the excluded rat, the LFP signals were particularly noisy, leading to an atypically low fraction of theta cycles categorized as periods of significant theta oscillation as defined below (0.38±0.02, compared to an average of 0.81 ± 0.06 for the other animals).

#### Tracking data

The position of the rat was determined as the midpoint between two LEDs mounted on its head. The 2D positions where then projected onto the linear track. We calculated instantaneous speed as the central difference of the position along the track, smoothed with a Gaussian filter with a standard deviation of 100 ms. Periods where rats ran continuously in the same direction from corner to corner were identified as rightward or leftward runs. A characteristic speed was calculated for each running direction in 4 cm spatial bins by taking the mean of speeds after discarding speeds below 10 cm/s that did not occur within 40 cm of either end of the track (e.g., see Figure 6B). This was done to avoid contamination of the calculation of the mean speed by infrequent cases where rats stopped in the middle of the track. Near the ends of the tracks, however, low speeds are the norm and therefore we do not set a minimum threshold there. A mean acceleration value was also calculated for each spatial bin by taking the difference on the smoothed speeds.

#### Firing rates

For calculating the firing rates used for identifying place fields and performing the Bayesian decoding, we discarded spikes emitted at running speeds below 10 cm/s that did not occur within 40 cm of either end of the track. This removed spikes produced on the rare occasions the animals stopped in the middle of the track. Virtually the same results were produced without this restriction (not shown). Firing rates were then calculated in 4 cm spatial bins and smoothed with a Gaussian filter with a standard deviation of 6 cm.

#### Identification of place fields

Candidate place fields were required to have a peak firing rate above 2 Hz and at least 25 spikes. The spatial extent of the field was identified by finding the positions at which the firing rate fell below 15% of the peak firing rate. If a place field was cut on either side by the end of the track, but the firing rate had gone below 66% of the peak firing rate before then, the field was accepted but marked as incomplete. If there were more than 3 consecutive spatial bins with zero occupancy between the field’s peak and one of the points at which the firing rate fell below threshold, the field was also marked as incomplete. Incomplete fields were excluded from the phase precession and place field skew analyses, and were treated differently in the place field size analysis (see below). Candidate place fields were screened manually to remove putative cells with overlapping phase precession clouds.

#### Place field sizes

For complete fields, place field size was defined as the distance between the points for which the firing rate fell below 15% of the peak firing rate. For incomplete fields, place field size was calculated as twice the distance between the field’s peak and the point on either side of the peak for which the firing rate fell below the 15% threshold. This ensured that a sufficient number of place fields at the ends of the track entered the analysis. Due to the pattern of place field skews along linear tracks (Diba and Buzsáki, 2008), we expect this method to slightly overestimate actual sizes, and thus the smaller place field sizes that we observed at the ends of the track should not be an artifact of the procedure.

For calculating the effect of running speed on place field sizes, we recalculated the firing rates with the same spatial bin size and smoothing as described above, but using data corresponding to periods with running speeds belonging to overlapping 20 cm/s wide speed bins in 10 cm/s steps starting at 2 cm/s. Rats spent too little time at some positions and speeds, often leading to an apparent firing rate of 0 due to the lack of time to produce any spikes. Thus, we defined a sampling index that quantifies whether the area where we expect the field to be (the extent of the place field as defined above) has been sampled well enough and we only attempt to measure the place field size in such cases. To obtain this sampling index, we summed the distances between all pairs of spatial bins within the field with occupancy above 300 ms, and divided it by the sum of distances between all pairs of bins in the field. This gives a number that increases nonlinearly from 0 (no valid bins) to 1 (all bins are valid) taking into account both the number of valid bins and how spread out they are through the field. We set the threshold for this measure at 0.4, which guarantees a good coverage. Place fields where then identified and their sizes calculated as described above. For the summary analyses, we removed extreme outliers below *Q*1 −*k· IQR* or above *Q*3 + *k · IQR* where Q1 and Q3 are the 25% and 75% percentiles, *IQR* = *Q*3 − *Q*1 and *k* = 3.

#### Phase precession slopes

Phase precession slopes were measured from linear fits of phase precession clouds falling within the extent of place fields as defined above. The fitting of phase precession clouds is complicated by to two matters: (i) phase is a circular variable, and (ii) the slope of the precession might be very steep. One possible way of dealing with the circularity of phase is to directly minimize angular distances between points and the best fitting line in the cylinder defined by position and phase. This can be done using circular-linear regression (Kempter et al., 2012; Schmidt et al., 2009) or by exhaustive search (O’Keefe and Recce, 1993). However, in such cases one must set the minimum and maximum slopes allowed, otherwise the optimal solution would be a line with a slope approaching ±∞ that loops around the cylinder going through all points. This would preclude us from correctly fitting clouds that really are very steep. Another possibility is to try to unwrap the phase precession cloud so that it lays uninterrupted in a plane and then perform a standard linear regression (Lenck-Santini and Holmes, 2008). However, this does not work well either for very steep phase precession clouds, since linear regression minimizes only the sum of square errors in phase, leading to excessively shallow fits of steep clouds.

We therefore developed a new method to determine phase precession slopes that works robustly even when slopes are very steep. It is based on the orthogonal distance regression, which minimizes the orthogonal distance of the data points to the best fitting line. This method introduces fewer *a priori* assumptions than standard linear regression, since it does not take position and phase to be the independent and dependent variables, respectively. Phase and positions were normalized to a range of [0, 1] to make their contribution equivalent in the calculation of orthogonal errors.

As commonly observed, spikes in the phase precession plot (theta phase vs. position) did not usually form a monotonic relationship. Instead, phase precession clouds slightly wrapped around 0° or 360°. This means that the theta phase, as extracted from the filtered LFP, does not accurately mark the transitions between actual theta cycles. Since we later analyze theta sequences, it is crucial to identify the proper beginning and end of the theta cycles. We did this by identifying the theta phase offset that optimized the phase precession relationship across all fields for each experimental session as follows. We considered potential offsets from −60° to 60° in 2° steps. For each value, we calculated orthogonal fits to the phase precession clouds for all fields, and then selected the phase shift that led to the lowest mean orthogonal error across all fields. The optimal shift corresponds to the phase shift for which the most points fall along uninterrupted phase precession clouds.

In addition to the ambiguity with respect to setting the boundaries of the theta cycle, for each spike that is fired near a boundary there is an ambiguity with respect to which cycle that spike should be attributed to. This could lead to large errors in the orthogonal fit and therefore distort the estimate of the precession slope. Thus, we modified the orthogonal fit for each field. We computed the slope and intercept of the line that minimized the sum of squared orthogonal distances of the points to the line with the following caveat. For points with normalized phases below 0.3, the orthogonal distance to the line that was minimized was either the distance of the original point, or of the point with its phase shifted by 1, whichever was smallest. Likewise, for points with normalized phases above 0.7 the smallest distances of either the original points or the points with their phases shifted by −1 were minimized. The minimization was done using the Nelder-Mead algorithm (Gao and Han, 2012) implemented in the SciPy library (Virtanen et al., 2020).

For calculating the effect of running speed on phase precession we proceeded as described for the place field sizes. We only attempted to calculate phase precession slopes when there were 12 or more spikes. Additionally, we analyzed phase precession slopes in single passes through a field. We calculated the average speed of the pass through the field from the tracking data. We focused on passes with a running speed above 2 cm/s, 6 or more spikes, a duration between the first and last emitted spikes above 400 ms and a coefficient of variation (standard deviation / mean) of instantaneous speed below 0.3. Passes with too many invalid positions in the tracking data where also excluded (sampling index < 0.4). For the summary analyses, we removed extreme outliers as described for place field sizes with *k* = 3 for phase precession slopes in the pooled analysis and *k* = 6 for single pass phase precession slopes.

#### Bayesian decoding and theta trajectory lengths

We performed a Bayesian decoding of position based on the ensemble spikes (Zhang et al., 1998) assuming a prior distribution over positions as in Davidson et al. (2009). The probability of the rat being at position *x* given a vector of spike counts for each cell, *n*, is given by

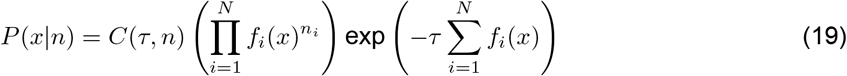

where *C*(*τ, n*) is a normalization factor which can be determined by the normalization condition ∑_*x*_ *P* (*x | n*) = 1 and *f_i_* is the firing rate of cell *i* in the corresponding running direction. We shifted theta phases so as to minimize the wrapping of phase precession clouds (see above), therefore we expected theta trajectories to be centered within individual theta cycles. Thus, we decoded within each cycle in overlapping 90° windows with a step size of 30° (no windows crossed theta cycles), thus *τ* changed slightly from cycle to cycle. The large window size was due to the need to gather a sufficient number of spikes for the decoding. We attempted to decode position only in phase bins where at least one spike was emitted, and for positions bins within a spatial extent of 70 cm centered at the current position. This value is below the average place field size at all running speeds (Figure 4C) and should therefore encompass the whole extent of the sweep.

When averaging theta cycles with similar speeds, we attempted to calculate theta trajectory lengths only when more than 5 cycles have been averaged.

When analyzing theta trajectories in individual cycles, we focused on cycles well covered by phase bins with peak decoded probabilities larger than 0.1. In particular, we required at least 5 such phase bins spanning a minimum of 210°.

Theta trajectory lengths were derived from a best fitting line. In order to calculate this fitting line, we first found the line that maximized the sum of decoded probabilities for positions within ±15 cm of the line (Davidson et al., 2009) and then performed linear regression with the points in this region weighted by their probabilities.

#### Place field skew

Place field skew was calculated as the ratio of the third moment of the place field firing rate distribution divided by the cube of the standard deviation (Mehta et al., 2000; Davoudi and Foster, 2019):

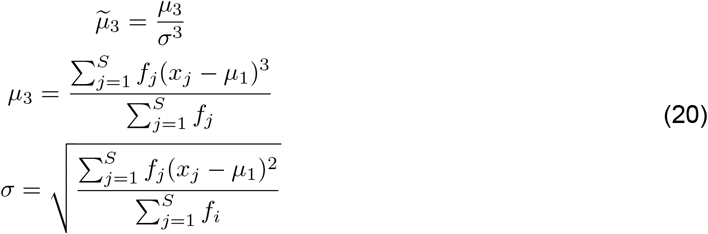

where, unlike above, *f_j_* is the firing rate of a given cell in the spatial bin *x_j_*, *S* is the number of spatial bins, and *μ*_1_ is the center of mass of the field:

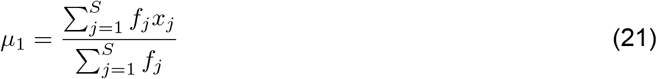

### Computational models

We generated surrogate data sets by taking the tracking and LFP data from experimental sessions and substituting the experimental spikes by those generated by our models under different coding assumptions. We discretized the session into 2 ms time bins and calculated, for each of them, the position that the cell population represents, *r*(*t*), according to some coding scheme and based on the phase of theta, *θ*(*t*). For the temporal sweep model, *r*(*t*) = *x*(*t* + *τ_θ_*(*θ*(*t*) − 180)/360), where *x*(*t*) is the position obtained from the tracking and *τ_θ_* is the time interval swept over in each theta cycle and which was set to 0.55 s in order to fit the experimental values for theta trajectory lengths. For the spatial sweep model, *r*(*t*) = *x*(*t*)+ *d_θ_*(*θ*(*t*) − 180)/360, where *d_θ_* is the spatial distance swept over in each theta cycle and which we set to 30 cm based on the fit to the theta trajectory lengths. Then, for each time bin, we generated spikes for each of 20 simulated place cells uniformly distributed along the track. Spikes were generated based on the activation level of each cell *i* at the represented position, as determined by the cells’ true place field. These true place fields were defined as Gaussian activation functions, *a_i_*, with a peak value of 1, discretized in 1 cm steps. The standard deviation of the Gaussian controls how tight the relationship is between firing at a particular phase and the time or distance to the peak’s center. A small standard deviation leads to very crisp phase precession clouds whereas a bigger one results in wider clouds with a noisier relationship between phase and position. For the temporal and spatial sweep models, the standard deviation was set to 7 cm, which produces phase precession clouds approximately matching the experimental ones.

Spikes were generated from a Poisson process with probability equal to an instantaneous firing rate times the length of the time bin:

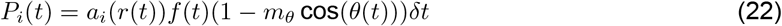

*f* (*t*) is the maximum allowed firing rate and was calculated as *f* (*t*) = 15 + 0.2*v*(*t*), where *v*(*t*) is the instantaneous running speed in cm/s. These values were chosen to approximate the increase in peak firing rate with running speed observed in the experimental data (Figure S4B). The term (1 − *m_θ_* cos(*θ*(*t*)) introduces an inhibitory modulation by the amplitude of the theta oscillation with *m_θ_* = 0.35 approximating the experimentally observed value in the experimental data set (not shown).

The behavior-dependent sweep model was the same as the spatial sweep model with two exceptions. First, the theta distance varies along the track being proportional to the average speed at each location. Second, we assume that the population code is noisy with different units coding for slightly different positions. Thus, we calculate *d_θ_*(*t*) and *r*(*t*) for each cell as:

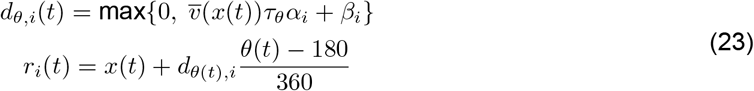

where *τ_θ_* = 0.57 s and 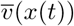 is the characteristic running speed from the tracking data at the current position. *α_i_* and *β_i_* represent multiplicative and additive noise terms respectively, and are derived from Gaussian distributions *N* (1, 0.2) and *N* (0, 10) and fixed for each cell for the entire simulation. As opposed the previous models, the width of the true fields was heterogeneous, and was set to min(4, 0.3*d_θ,i_*).

## Acknowledgements

We thank Loren Frank for insightful discussions and Laia Serratosa Capdevila for feedback on the draft. This work was supported by grants from the German Research Foundation (DFG), grant 419037518 – FOR 2812, P2 (S.C.), from the German Federal Ministry of Education and Research (BMBF), grant 01GQ1506 (S.C.), and from the National Institute of Mental Health, grant NIMH R01MH109170 (K.D.).

## Author Contributions

Conceptualization, E.P., S.C.; Methodology, E.P., K.D., S.C.; Software, E.P.; Formal Analysis, E.P., K.D., S.C.; Writing – Original Draft, E.P., S.C.; Writing – Review & Editing, E.P., K.D., S.C.; Visualization, E.P.; Supervision, S.C.; Project Administration, S.C.; Funding Acquisition, K.D., S.C.

## Declaration of interests

The authors declare no competing interests.

## Appendix: Variable theta phase locking fails to account for the combination of population and single-cell results

For completeness, we also considered whether the experimental results could be accounted for by a fourth model similar to the one proposed by Chadwick et al. (2015). Cells participated in homogeneous spatial sweeps, but the coordination between cells increased with running speed. At lower speeds, the lack of coordination between cells should bias the decoder towards the current position, leading to shorter decoded theta trajectories.

We modeled this by introducing the following modifications to the behavior-dependent sweep. The theta distances were constant:

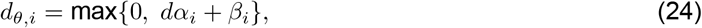

with *d* = 35. The standard deviation setting the width of the true place fields was min(3, 0.25*d_θ,i_*). To vary the coordination between cells with speed, we introduced into each cell independently a certain amount of theta phase noise that decreased with running speed:

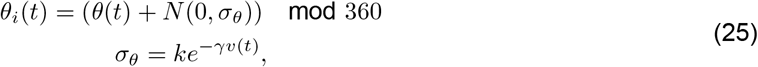

with *k* = 120 and *γ* = 0.025.

As expected, the model provides a good fit for the increase in theta trajectory lengths with running speed. However, the model did not account for the increase in pooled place field sizes and phase precession slopes without within-field changes (Figure S5A, B).

To confirm that the shorter decoded theta sequences at lower speeds are not caused by decreased coordination between pairs of cells, we further calculated the effect of speed on the variance in pairwise phase differences between spikes of cells with overlapping fields. First, we found cells whose place fields overlapped by more than 20 cm. Then, when rats were within 10 cm of the overlapping region of two cells, we collected, for each theta cycle individually, the phases at which spikes from each cell were emitted, and calculated all pairwise phase differences between them. Finally we calculated their circular variance (Fisher, 1993) as:

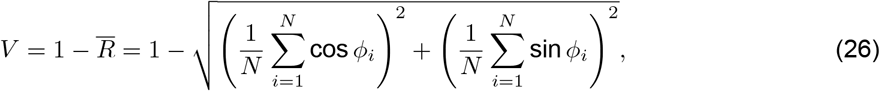

where *ϕ_i_* is *i*-th pairwise phase difference between the two cells.

If the cells code for positions in a coordinated fashion, the variance in phase differences between their spikes should be low. We found that the level of variance present in the variable noise model at low speeds was unrealistically high as compared to the experimental data (Figure S5C-E). Furthermore, in the experimental data, the variance in pairwise phase differences for a cell pair did not significantly change with running speed for any of the animals (*p* > 0.15; Wilcoxon signed-rank test). Overall, we conclude that these results rule out the possibility that changes in cell coordination at the theta time scale play a major role in explaining the changes observed in theta trajectory lengths.

## Supplementary Figures

**Figure S1:**
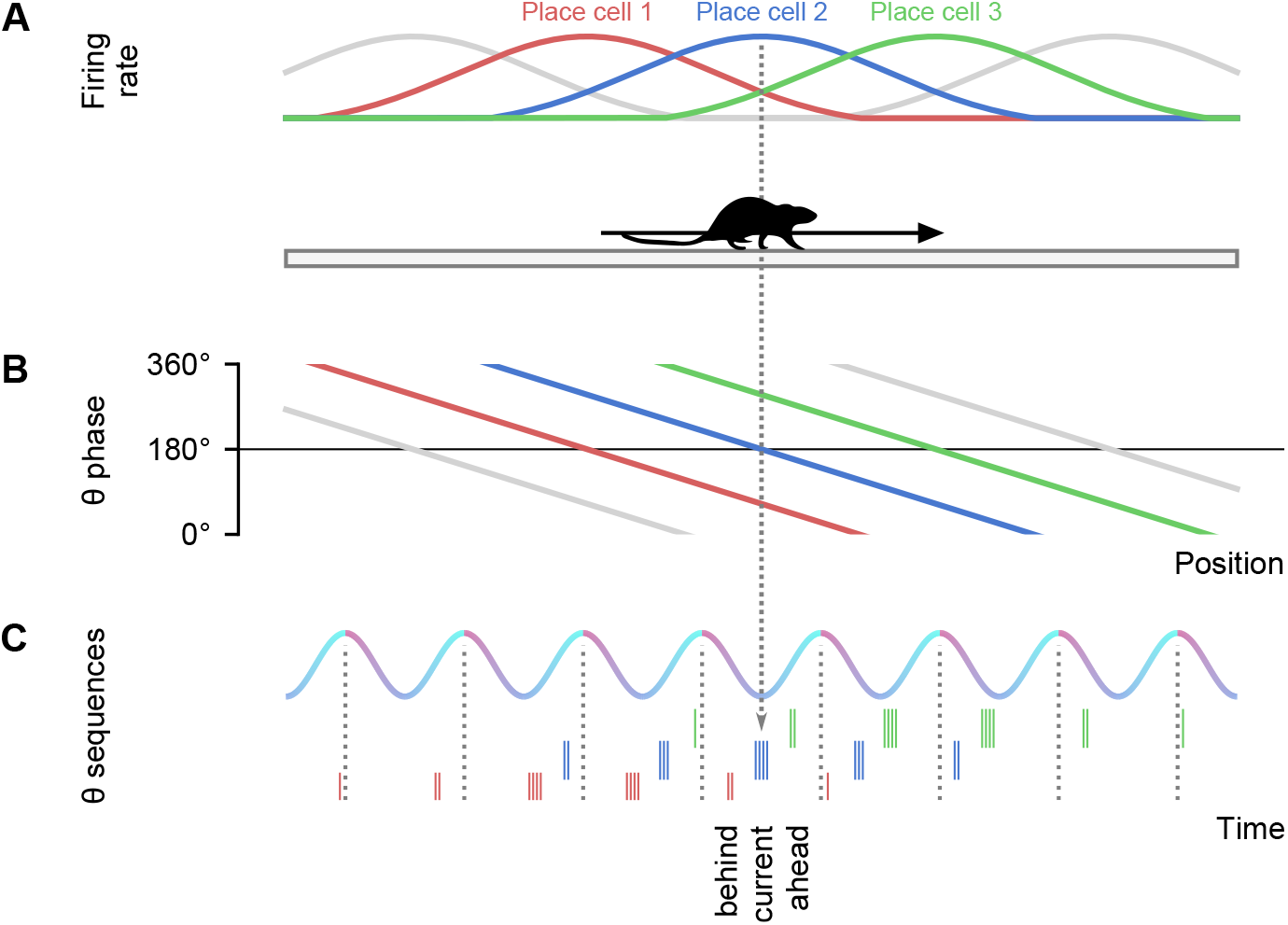
Illustration of theta phase coding in spatial navigation. Related to Figure 1 and 2. **A**: A rat is running from left to right on a linear track. The firing rate of three place cells are indicated in different colors. **B**: The idealized theta phase vs. position relationship for the spikes emitted by each of the cells shows a decrease in phase as the animal crosses the field (theta phase precession). **C**: At the population level, phase precession manifests as spike sequences that represent, in a temporally compressed fashion, the sequence of place fields being traversed. The falling edge of the oscillation holds spikes from place cells with fields centered behind the current position of the rat, followed by spikes from cells with fields centered at the current position of the rat at the trough of the oscillation, and place cells with fields centered ahead in the rising edge.

**Figure S2:**
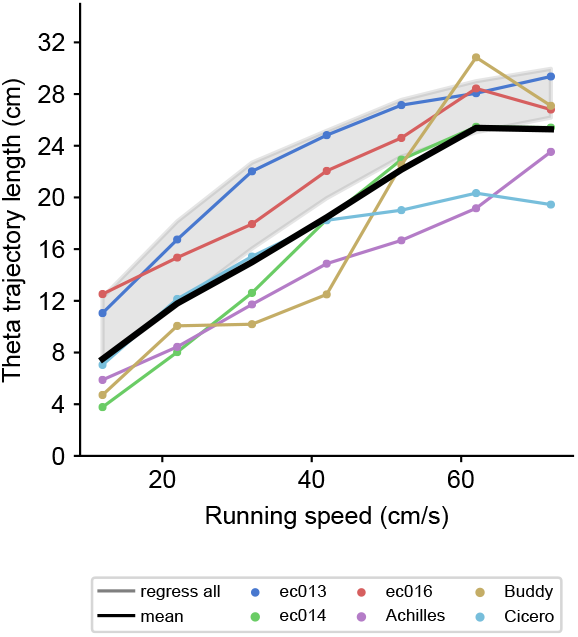
Theta trajectory lengths remain unchanged when restricting the analysis to theta cycles occurring in areas covered by place field analyses. Related to Figure 3. Same analysis as in Figure 3D, but pooling only theta cycles occurring in portions of the track that are covered by place fields included in the place field size or phase precession slope analyses.

**Figure S3:**
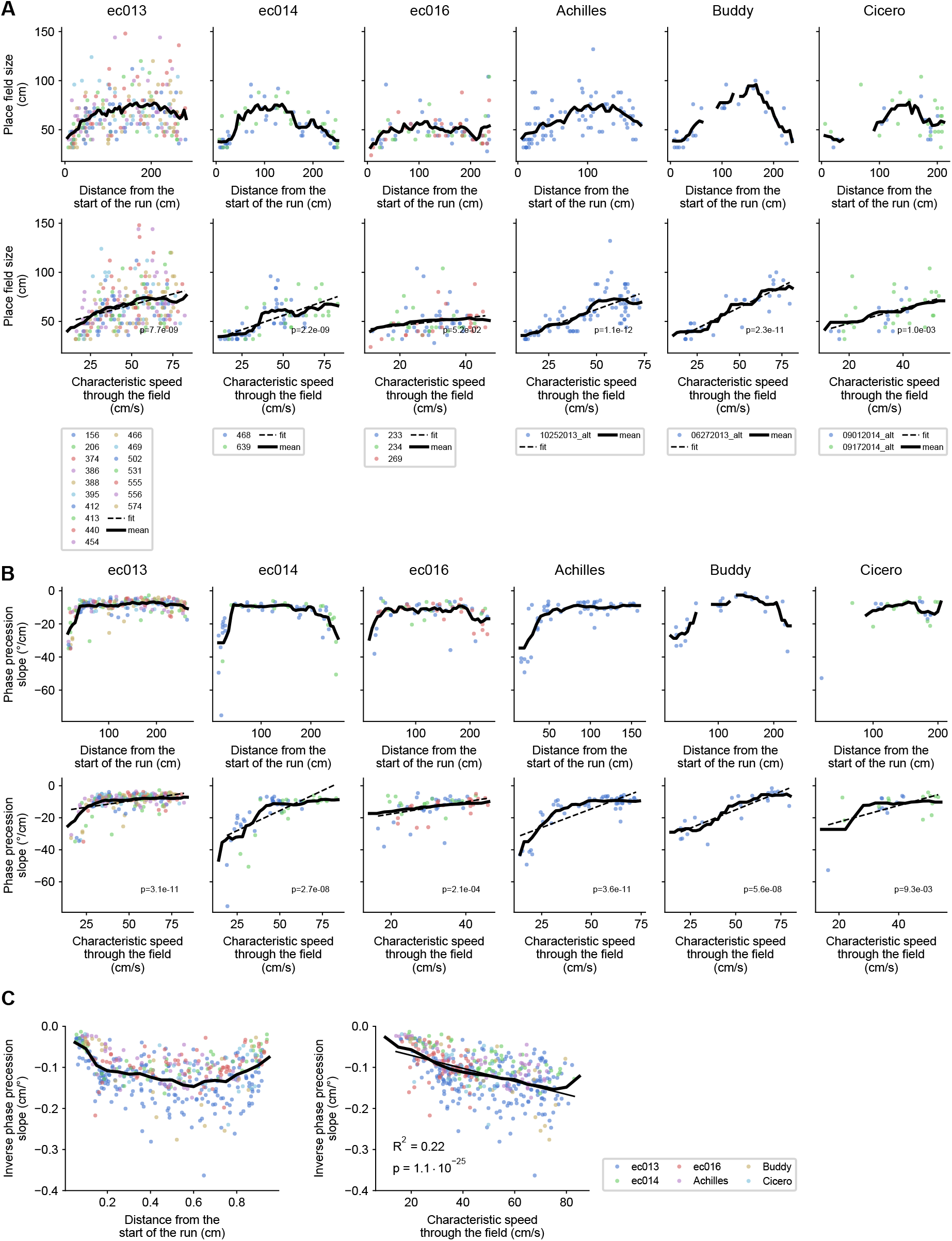
Place field sizes and phase precession slopes increase with characteristic running speed. Related to Figure 6. **A**: Place field sizes as a function of distance from the start of the run (first row), and characteristic speed through the field (second row). Each column contains data for one animal. Different colors for each animal represent different sessions. Each point is a place field. Dashed lines indicate linear fits (with their associated p values; Wald Test with t-distribution of the test statistic), and thick black lines indicate moving averages with sliding windows of 30 cm and 20 cm/s for distance and speed, respectively. **B**: Same as above but for phase precession slopes. **C**: The inverse of phase precession slope pooled across animals plotted against normalized run distance or characteristic speed through the field. Thick black line represents the moving average on sliding windows of 0.1 and 15 cm/s for the normalized distances and speed, respectively.

**Figure S4:**
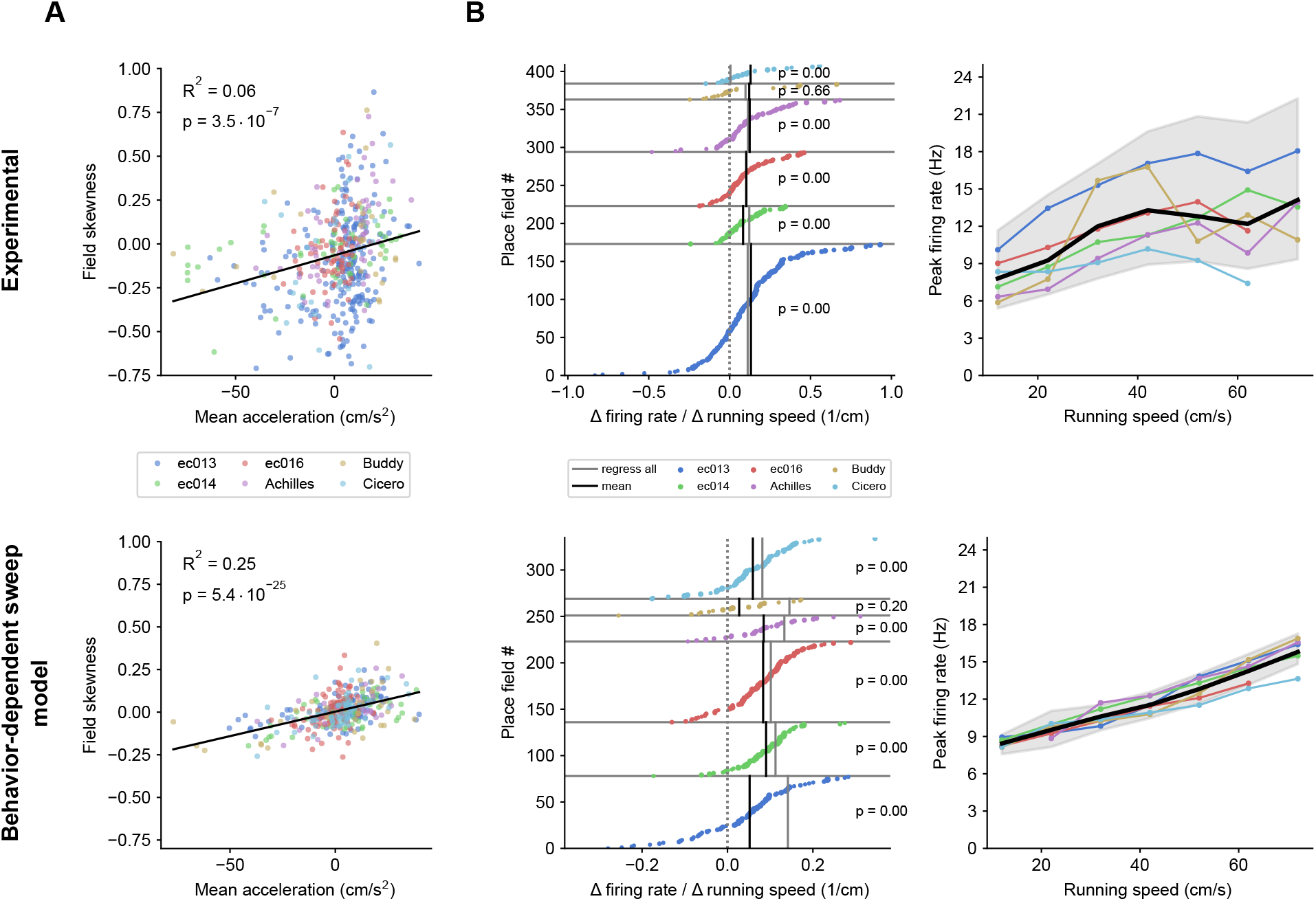
The behavior-dependent sweep model captures changes in place field skew and firing rates. Related to Figure 7. **A**: Place field skewness is significantly correlated with the mean acceleration through the field in the experimental data (top row) and the model (bottom row), although the experimental distribution is notably broader. Black lines represent linear fits. P values for the fits are indicated (Wald Test with t-distribution of the test statistic). **B**: The behavior-dependent sweep model replicates the experimentally observed increase in firing rates with running speed. Analyses and plotting conventions as for Figure 4B, C.

**Figure S5:**
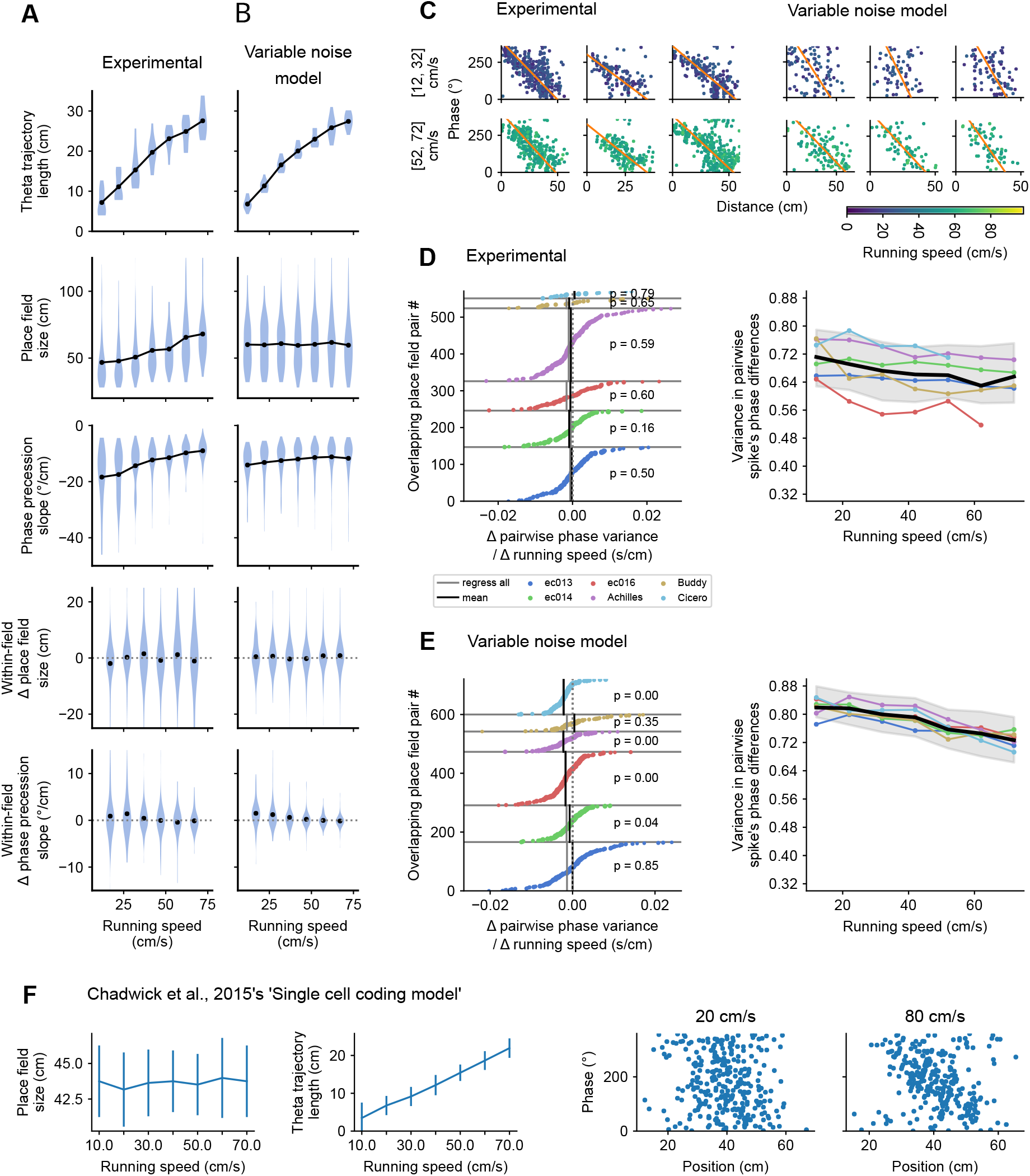
Variable theta phase locking fails to account for the combination of population and single-cell results. Related to Figure 7. **A, B**: Comparison of experimental and model results. The variable noise model does a good job at capturing the increase in theta trajectory lengths with running speed, but it does so at the cost of within-field changes in phase precession slopes at lower speeds and without capturing any of the increase in pooled place field sizes. **C**: Example phase precession clouds at low and high speeds for experimental and modeled place fields. Each column corresponds to a cell. The variable noise model introduces an atypically large amount of noise at low speeds as compared to the experimental data. **D, E**: Analyses and plotting conventions as for Figure 4B, C, but for the circular variance in pairwise phase differences between spikes emitted by cells with overlapping place fields. Note that the variable noise model produces atypically high variance and the variance tends to reduce with speed for individual cell pairs, whereas this effect is not significant in the experimental data. Continues on the next page. **F**: A reproduction of the model by Chadwick et al. (2015) which also operates by increasing phase locking with running speed. With the parameters published in their study we can indeed reproduce the increase in theta trajectory lengths with running speed reported by Maurer et al. (2012) (left). However, like in our variable noise model, place field sizes remain constant with running speed (middle) and phase precession clouds look unrealistically noisy at lower speeds (right). The two panels on the right display the phase precession clouds for one cell at low and high running speeds (all cells in the model behave identically). We ran their model on 30 sessions composed of 20 idealized laps through a track at constant speed. Error bars represent standard deviations.

## Notes

### Competing Interest Statement

The authors have declared no competing interest.

